# Deep mutational scanning of whole SARS-CoV-2 spike in an inverted infection system

**DOI:** 10.1101/2023.07.17.549430

**Authors:** Shunta Taminishi, Songling Li, Yusuke Higuchi, Yuhei Kirita, Daisuke Motooka, Yuki Ozaki, Takao Arimori, Nariko Ikemura, Yumi Itoh, Satoaki Matoba, Toru Okamoto, Junichi Takagi, Daron M Standley, Atsushi Hoshino

## Abstract

In order to investigate SARS-CoV-2 mutations and their impact on immune evasion and infectivity, we developed a Deep Mutational Scanning (DMS) platform utilizing an inverted infection assay to measure spike expression, ACE2 affinity, and viral infectivity in human cells. Surprisingly, our analysis reveals that spike protein expression, rather than ACE2 affinity, is the primary factor affecting viral infectivity and correlated with SARS-CoV-2 evolution. Notably, within the N-terminal domain (NTD), spike expression and infectivity-enhancing mutations are concentrated in flexible loops. We also observed that Omicron variants BA.1 and BA.2 exhibit immune evasion through receptor binding domain (RBD) mutations, although these mutations reduce structural stability. Interestingly, the NTD has evolved to increase stability, compensating for the RBD instability and resulting in heightened overall infectivity. Our findings, available in SpikeScanDB, emphasize the importance of spike expression levels and compensatory mutations in both the NTD and RBD domains for shaping Omicron variant infectivity.

## Introduction

Since the emergence of SARS-COV-2, the spike gene has continued to evolve under the selection pressure of better transmissibility and immune evasion. The D614G substitution that appeared early in the COVID-19 pandemic shifted the receptor binding domain (RBD) conformation toward an open state to better bind an entry receptor, ACE2(Korber et al., 2020; Yurkovetskiy et al., 2020). The P681R mutation, highly conserved in Delta lineages, facilitated cleavage of the spike protein and enhanced viral fusogenicity and replication, which resulted in higher transmission and pathogenicity(Mlcochova et al., 2021; Saito et al., 2022). In addition to transmission fitness, SARS-CoV-2 variants have acquired resistance to neutralization from vaccinated and convalescent sera as well as therapeutic antibodies. The Beta and Mu strains exhibited some degree of immune evasion from serum neutralizing antibodies(Uriu et al., 2021). After the Delta wave, the Omicron variant emerged with substantial mutations compared to any previously described SARS-CoV-2 isolates, comprising 26 to 32 amino acid level changes in the spike protein. Among these,15 mutations are located in the RBD and induced striking immune evasion(Ikemura et al., 2022).

Deep mutational scanning (DMS), which consists of saturation mutagenesis and deep sequencing analysis, is a powerful tool to evaluate the functional alterations in a protein due to amino acid substitutions. DMS has previously been conducted to understand the mutational effects of SARS-CoV-2 spike; however, of these studies employed yeast surface display of the RBD, not full-length spike, which limited the scope to RBD expression, ACE2-binding and antibody escape(Greaney et al., 2021; Starr et al., 2021a; Starr et al., 2021b; Starr et al., 2020). Therefore, the functional roles of other domains have remained largely unknown. Unlike previous studies, we employed a human cell expression system to fully express the entire spike protein and accurately replicate the process of SARS-CoV-2 infection using ACE2-coating pseudovirus. In this system, the spike protein interacted with ACE2 and triggered membrane fusion between the cell and the virus, albeit in an inverted orientation (Ikemura *et al*., 2022). This approach enabled the direct evaluation of infectivity and an understanding of its relationship with spike expression and binding affinity toward ACE2. The target region could also be expanded to the whole spike, including the N-terminal domain (NTD) and the S1/S2 junction.

Through extensive DMS analysis, we discovered that heightened infectivity was primarily linked to elevated levels of spike protein expression, while enhanced receptor binding domain (RBD) affinity towards ACE2 had a relatively smaller impact on viral infection. The spike RBD is the main target of neutralizing antibodies from vaccinated and convalescent sera, but the anti-NTD antibodies can also moderately neutralize infection(Cerutti et al., 2021; Ikemura *et al*., 2022; McCallum et al., 2021). Within the NTD, three loops--N1 (residues 14-26), N3 (residues 141–156) and N5 (residues 246–260)--form an antigenic supersite and NTD-targeting antibodies are susceptible to mutations in these loops(Cerutti *et al*., 2021; McCallum *et al*., 2021). Functionally, our DMS analysis further unveiled an accumulation of mutations that enhance spike expression and infectivity specifically within flexible loops positioned at the periphery of the spike protein. Domain-wise analysis of Omicron BA.1 and BA.2 spikes indicated that the Omicron RBD lost structural stability in exchange for a gain in immune evasion, but that NTD mutations compensated for the fragile Omicron RBD in order to preserve infectivity. When BA.1 NTD mutations were categorized into 4 clusters, only G142D/Δ143-145 contributed to immune evasion consistent with previous NTD antigenicity analysis. The additional mutations, A67V/Δ69-70, T95I, and Δ211/L212I/ins214EPE, were involved in the preservation of spike expression and infectivity. DMS in the inverted infection assay thus provided comprehensive insights into functional alterations upon spike mutation, which has further deepened our understanding of viral evolution.

## Results

### Deep mutational scanning in an inversely oriented infection assay

To understand the effect of SARS-CoV-2 spike mutations on its characteristics, including protein expression, binding-affinity toward ACE2 and consequent viral infectivity, we conducted deep mutational scanning of the whole spike in the context of full-length spike protein expression in human Expi293F cells(Ikemura *et al*., 2022). Original signal sequence was replaced with HMM38 synthetic one(Barash et al., 2002) and HA tag was inserted in the N-terminal of the spike. Assay was independently conducted in each domain and some restriction sites were introduced between the domains for library construction (Figure 1A). The HA-tagged spike library encompassing all 20–amino acid substitutions based on the ancestral SARS-CoV-2 Wuhan strain was transfected into Expi293F cells under conditions where cells in general expressed no more than a single variant(Chan et al., 2021) (Figure 1B). The spike expression level was determined by anti-HA tag antibody and the binding affinity was evaluated by two-dimensional analysis with anti-HA tag antibody and ACE2-Fc chimera protein. For spike expression analysis, the top and bottom 25% of cells among HA-positive population were collected by fluorescence-activated cell sorting (FACS) (Figure 1C). Similarly, the highest and lowest 25% of cells in terms of the ACE2-Fc signal were gated in the distribution with spike expression level to analyze the binding affinity (Figure 1D). The DMS library expressing cells were also incubated with ACE2-coating GFP reporter pseudoviruses. This infection assay functionally reproduced the process of membrane fusion with spike and ACE2 between viruses and host cells, even though it was in an inverse orientation (Figure 1E). This inverted infection system enabled sequencing of the spike mutants in a large-scale pool library. In the RNA-based deep sequencing of DMS library, the read count ratio of two populations was normalized by the silent mutation construct and log 10 scaling was used to define the spike expression, affinity, or infectivity value. To validate the accuracy of the inverted infection assay, we compared it with the SARS-CoV-2 pseudovirus infection assay and observed high correlation of infection rate in major SARS-CoV-2 variants (Fig. 1F). GFP-positive cells with ACE2-coating pseudoviurs infection and GFP-negative control cells were harvested and log 10 scaling of the read count ratio was similarly defined as infectivity value.

**Figure 1.**
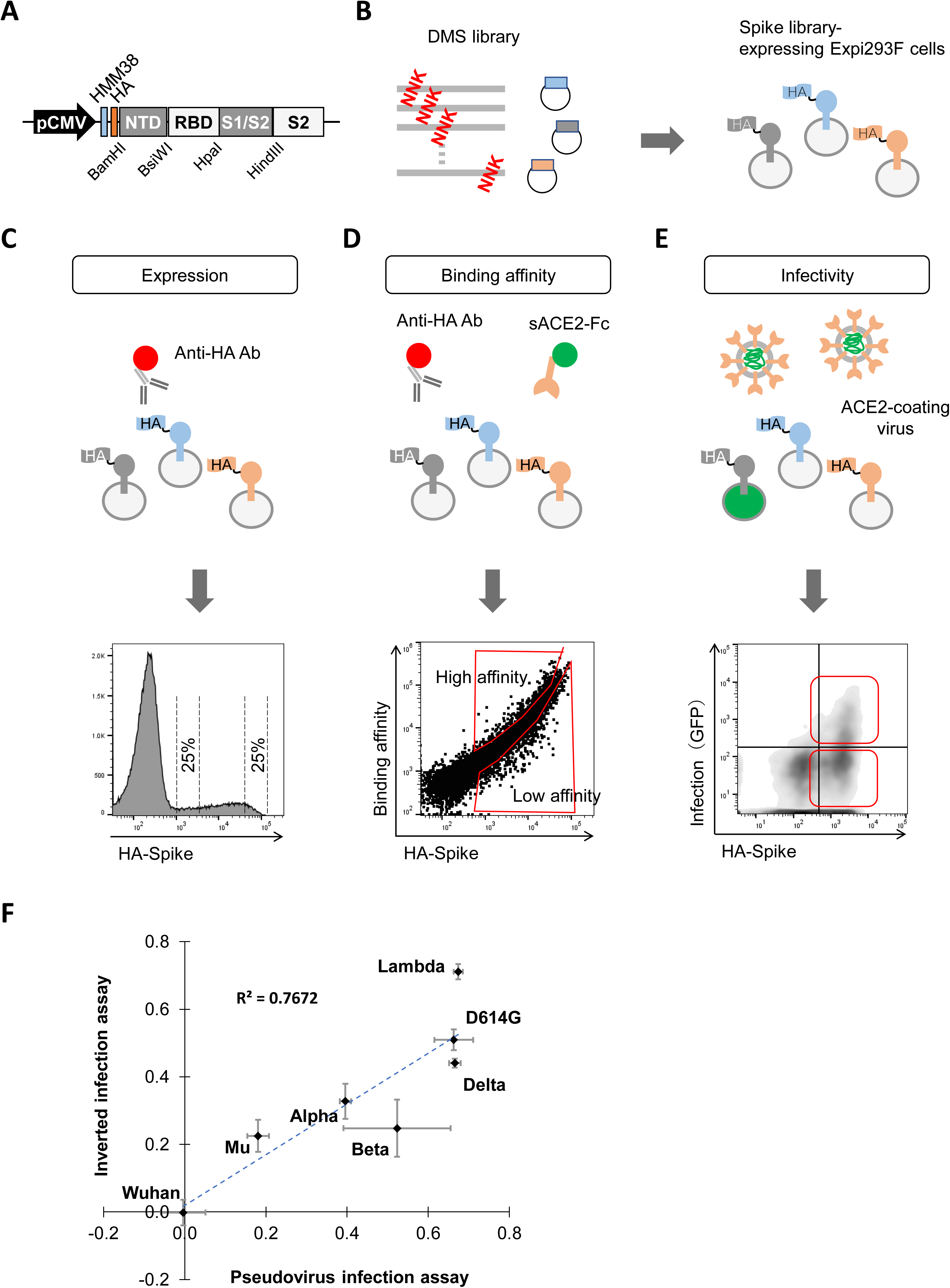
Deep mutational scanning in the inverted infection assay. (A) A design of the deep mutational scanning (DMS) library for the spike of SARS-CoV-2. HA tag was introduced in the N-terminus of the spike. Restriction sites were also introduced between the domains to insert library fragments. (B-E) Schematic of deep mutational scanning (DMS). DMS library was developed using pooled synthesized DNS with NNK mixed bases and expressed Expi293F cells in the context of full-length spike (B). Spike expression level was analyzed with the gating of the highest and lowest 25% HA signal among HA-positive cells C. For affinity analysis, library expressing cells were incubated with anti-HA antibody and ACE2-Fc. The highest and lowest 25% of cells with ACE2-Fc binding relative to HA-spike expression were collected (D). Infectivity was examined in the assay where DMS library expressing cells were incubated with ACE2-coating GFP reporter pseudovirus. GFP-positive and negative cells was sorted (E). (F) Correlation between luciferase-reporter SARS-CoV-2 pseudoviruses and inverted infection assay among major variants. Data are mean ± SEM from three biological replicates.

### DMS in the N-terminal domain

In addition to the RBD, SARS-CoV-2 variants frequently acquire mutations in the spike NTD. To comprehensively understand the functional importance of these mutations, we conducted DMS for the NTD. DMS experiments were performed in duplicate and produced similar results; the R2 (coefficient of determination) values were 0.6039, 0.8725, and 0.8589 for infectivity, expression, and binding affinity, respectively (Figure S1A). An overview of how amino acid mutations affected infectivity, expression and ACE2 binding affinity was visualized as heatmaps (Figure 2A). We next performed a series of experiments to confirm the accuracy of these DMS results using NTD mutations observed in major variants, including L18F and D80A in Beta, D138Y and R190S in Gamma, T19R and G142D in Delta, and T76I in Lambda. Infectivity was evaluated by luciferase reporter pseudovirus infection in ACE2-expressing HEK293T (HEK293T/ACE2) cells. The pseudovirus harbored a small luminescent peptide tag, HiBiT to precisely normalize input virus doses(Ozono et al., 2021). There was strong correlation (R2=0.8599) between DMS infectivity values and the log 10 scaled luciferase signal relative to that of parental virus. The individual mutant spike expression was determined by flow cytometry for HA-tagged spike protein. Similarly to infectivity, the log 10 scaled HA signal relative to that of parent virus was well correlated with the DMS value (R2=0.8914). The individual affinity was quantified as the ACE2-Fc signal in an arbitrary range of spike expression (Figure S1B). There was reasonable correlation between the log 10 scaled relative ACE2-Fc signal and DMS affinity (R2=0.5993) (Figure 2B). Together, these robust validations supported the quality of the DMS data.

**Figure 2.**
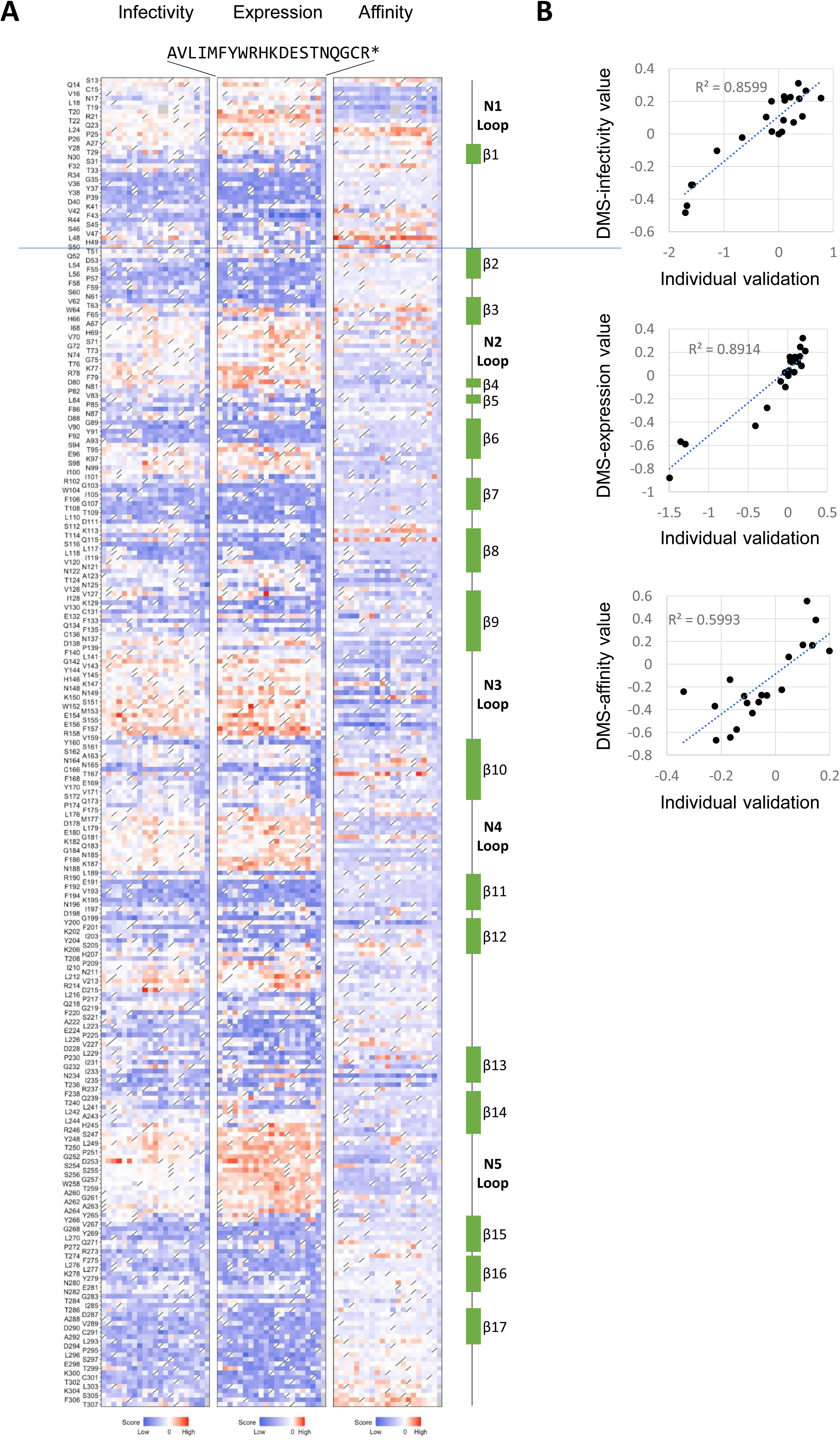
Deep mutational scanning for N-terminal domain. (A) Heatmap showing how all single mutations affect the infectivity, spike expression, and affinity toward ACE2 in the N-terminal domain. Squares are colored by mutational effect according to scale bars on the bottom, with blue indicating deleterious mutations. Squares with a diagonal line through them indicate the original Wuhan strain amino acid. (B) Validation of DMS results using NTD mutations including major variants ones. Infectivity was verified by the standard infection assay with luciferase reporter SARS-CoV-2 pseudovirus and HEK293T/ACE2 cells. Spike expression was examined by the same assay as DMS and the strategy of affinity assessment was explained in Figure S1B.

### Infectivity is dominantly determined by spike protein expression level

Heatmap visualization indicated a similar pattern in infectivity and spike expression (Figure 2A). Statistically, infectivity was substantially correlated with expression (R2=0.7206), but not with binding-affinity (R2=0.0052) (Figure 3A). Mapping to the spike trimer structure indicated that mutants with enhanced infectivity and spike expression accumulated in the five NTD-loop regions that are located at the periphery of the spike. In contrast, mutations enhancing binding affinity were enriched in the region facing the RBD or S2 (Figure 3B). Many of these mutations were associated with decreased protein expression and therefore were less infectious overall (Figure 2A). Despite the dominant impact of spike expression on infectivity, there were some exceptional residues where binding affinity determined the infectivity. Among N-glycosylation sites, mutations in N165/T167 increased the binding affinity and some amino acid substitutions resulted in higher infectivity (Figures 2A and 3B). Conversely, mutations in N234/T236 decreased the affinity and infectivity, even though they increased spike expression (Figures 2A and 3C). The structural and functional roles of spike protein glycosylation have been analyzed by structural studies and molecular dynamics (MD) simulations starting from the atomic structures(Casalino et al., 2020; Choi et al., 2021; Mori et al., 2021). When the RBD is closed, the N165 glycan is located above the RBD and N234 glycan is located below the RBD, where both glycans stabilize the closed conformation with frequent interaction with the closed-state RBD(Casalino *et al*., 2020; Choi *et al*., 2021; Mori *et al*., 2021). When the RBD is open, the high-mannose-type N234 glycan intrudes in a cavity between the RBD and the S2 subunit and forms hydrogen bonds with S2, reinforcing the open-state from below(Choi *et al*., 2021; Mori *et al*., 2021). Our DMS results were consistent with these mechanistic models, although one previous study showed that deletion of the N165 glycan reduced the binding affinity toward ACE2(Casalino *et al*., 2020). At sites unrelated to glycosylation, increased infectivity due to allosterically enhanced binding affinity was observed in L48, S50, K113, and Q115 (Figures 2A and 3B).

**Figure 3.**
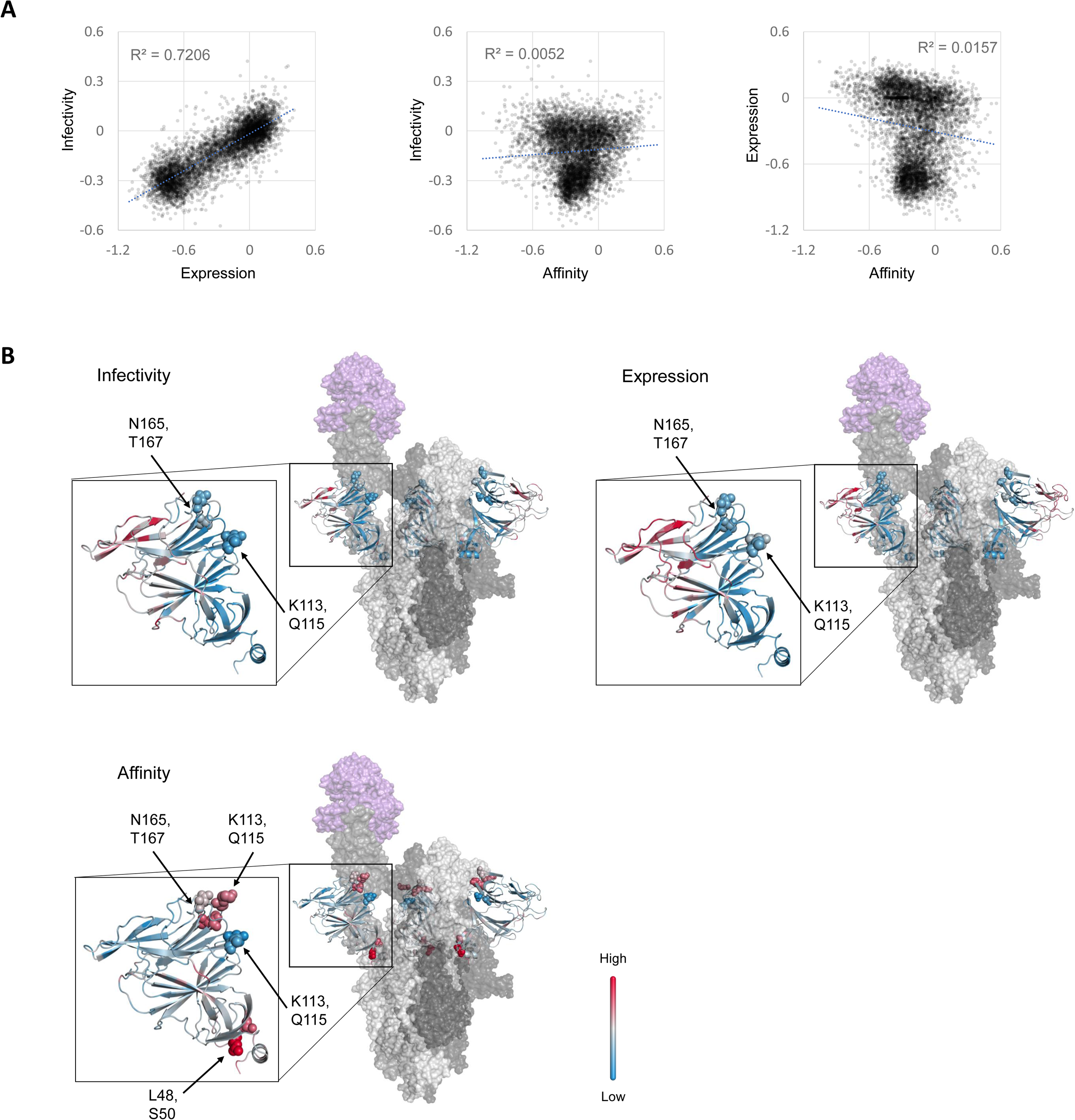
The spike expression mainly contributes to SARS-CoV-2 infectivity. (A) Correlation between functional alterations of infectivity, spike expression and affinity in DMS for the NTD. (B) Projection of functional changes due to mutations onto SARS-CoV-2 spike trimer with one RBD in the open state built via homology modelling. The values of infectivity, expression and affinity were summed up in each residue and indicated according to scale bars on the right. Sphere models are indicated residues in the results. ACE2 is colored in purple. Each spike monomer is in light, middle, and dark gray.

### Increased binding affinity contributes little to infectivity

DMS for affinity was also conducted in the RBD region and we analyzed this data together with infectivity and spike expression data that we previously reported(Ikemura *et al*., 2022) (Figure S2A). Similarly to DMS for the NTD, dominant involvement of spike expression in the infectivity was observed in the RBD (R2=0.712; Figure S2B). Previously, Starr et al. conducted DMS for the RBD and evaluated the total ACE2 binding, not affinity, as a surrogate for infectivity. In their data, ACE2 binding correlated well with expression (R2=0.4278; Figure S2C). To further examine the contribution of affinity-enhancing RBD-ACE2 interface mutations to infectivity, we prepared cells expressing the ACE2 mutant 3N39, which has a 100-fold higher affinity (Higuchi et al., 2021) and observed viral infectivity. As protein expression levels affect infectivity, HA tagged wild-type ACE2 and 3N39 were stably expressed in HEK293T cells with lentivirus and cells were sorted according to the arbitrary HA signal to equalize ACE2 expression between wild-type and 3N39 (Figure S3A). Authentic SARS-CoV-2 infection and replication were comparable between wild-type ACE2 and 3N39 expressing cells (Figure S3B). Similar infectivity was observed in a SARS-CoV-2 pseudovirus infection assay (Figure S3C). These results collectively emphasized the importance of spike protein expression levels in viral infectivity, as compared with the binding affinity toward ACE2.

### DMS in the S1/S2 junction

S1/S2 junction from F541 to S750 was analyzed in duplicate with high reproducibility (Figure S4A and B). D614G is the first predominant circulating vital mutant and increased infectivity and transmissibility was reported both in vitro and in vivo(Hou et al., 2020; Ozono *et al*., 2021). The D614G mutation breaks a salt bridge between D614 and K854 in the fusion peptide proximal region, which contributes to the RBD-up conformation(Cai et al., 2020). In addition, the disordered loop in the D614 S trimer wedges between domains within a protomer in the G614 spike. This interaction increases the number of functional spikes and enhances infectivity(Zhang et al., 2021). Consistent with this mechanism, our DMS results indicated that the D614G mutation enhanced affinity, spike expression, and consequent infectivity (Figure S4A). The dominant contribution of spike expression to infectivity was also observed in the S1/S2 junction (Figure S4C). Interestingly, affinity-enhancing mutations were frequently observed here unlike other domains and they tended to be negatively correlated with spike expression. One exception was the furin cleavage site, R682 to V687, where both affinity and spike expression were strikingly upregulated. Furin recognition site mutations resulted in increased affinity and stabilized the S1/S2 structure through the prevention of S1/S2 cleavage; However, infectivity was little affected. There are two pathways for SARS-CoV-2 infection. One is the S2′ site cleavage by TMPRSS2 and cellular entry (cell surface entry). The other is endocytosis and lysosomal cathepsin-mediated spike cleavage, which occurs when TMPRSS2 is unavailable (endosomal entry)(Jackson et al., 2022; Takeda, 2022). Furin-mediated S1/S2 cleavage promotes S1 shedding and exposes the S2′ site to TMPRSS2(Jackson *et al*., 2022). It means that the loss of furin-cleavage site impairs cell surface entry but consequent structural stability and increased affinity facilitate endosomal entry. Since the HEK293T cells we used here modestly express TMPRSS2(Partridge et al., 2021), the overall infectivity was nearly unchanged.

### Spike expression level is involved in SARS-CoV-2 evolution

We next examined the contribution of spike expression and ACE2-binding affinity to human SARS-CoV-2 evolution. Mutation frequency in naturally circulating SARS-CoV-2 was adopted from the previous report which included mutations expected to have ≥20 occurrences(Dadonaite et al., 2023). The results of DMS analyzing infectivity and spike expression were reasonably correlated with the enrichment of mutations observed in naturally circulating SARS-CoV-2, whereas DMS for affinity exhibited no association (Figures 4A-C). Consistent with the impact on infectivity, these results indicates that spike expression, not ACE2 binding affinity, is the predominant factor for SARS-CoV-2 evolution. This trend was observed similarly in NTD, RBD and S1/S2 junction (Figures S5A-C). Our DMS in the inverted infection assay was correlated with the previous DMS for pseudovirus infection (Figure 4D) and both DMS measurements comparably reflected the actual enrichment of SARS-CoV-2 mutations(Dadonaite *et al*., 2023). The DMS for infection reasonably correlated with viral evolution, but it was somewhat limited. The first reason is that not only infectivity but replication activity determines virus transmissibility. The P681R mutation is known to be a determinant for enhanced fusogenisity and viral replication fitness of the Delta strain(Liu et al., 2022; Saito *et al*., 2022); other spike mutations could similarly affect transmissibility through replication activity. The second is epistasis. It is known that effects of some mutations in Omicron spike RBD differ from those measured in the ancestral background. For instance, Q498R and N501Y substitutions contribute to ACE2 binding in Omicron with more impact than in earlier viral strains(Starr et al., 2022). Concomitant mutations could contribute nonadditively compared with single amino acid mutations. The last reason is that immune-evading mutations is a driving force for SARS-CoV-2 evolution and reduces the impact of infectivity on fitness in naturally circulating viruses.

**Figure 4.**
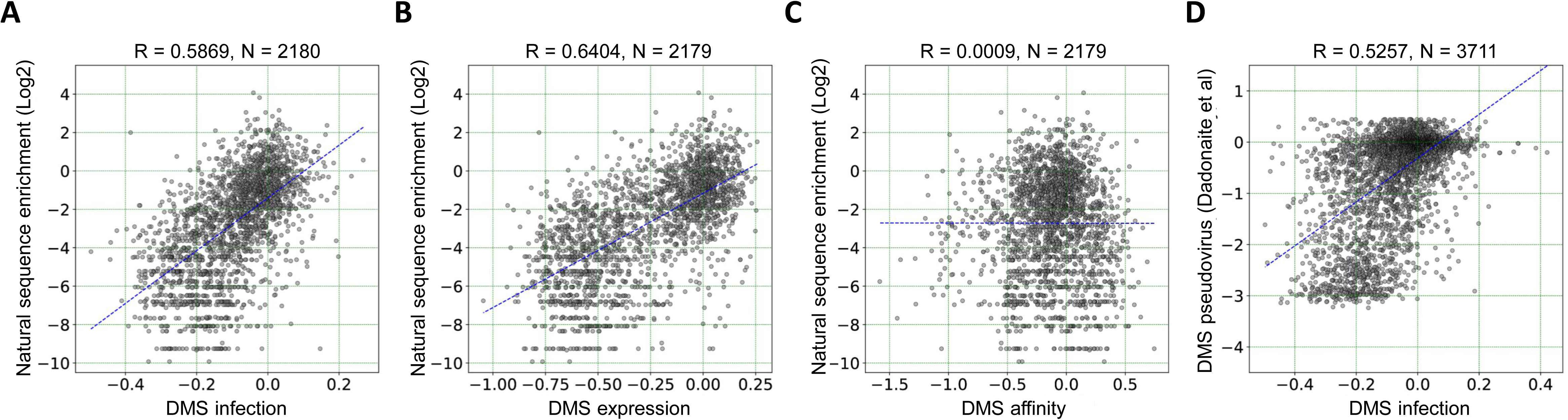
Functional effects of spike mutations on the viral transmission fitness. (A-C) Correlation between the mutation frequency of naturally circulating SARS-CoV-2 and functional alteration of spike including infectivity (A), spike expression (B), and affinity toward ACE2 (C). These correlations in each domain were presented in Figure S5. (D) Correlation between our infectivity DMS in the inverted infection assay and Dadonaite et al. pseudotyped lentivirus-mediated DMS(Dadonaite *et al*., 2023).

In position-wise analysis, previous DMS studies reported a correlation between relative accessible surface area (RSA) and spike expression-affecting mutations in the RBD(Ouyang et al., 2022; Starr *et al*., 2020). In general, solvent inaccessible buried residues are known to be important for proper protein folding and structural stability. Here, structural flexibility was further quantified by the change of RSA of each residue position in 607 trimeric spike complex structures collected from Protein Data Bank (PDB) (Bekker et al., 2022). Spike complexes were decomposed into monomeric chains, from which residue-wise RSA by the program NACCESS (Hubbard and Thornton, 1993) and standard deviation of RSA (dRSA) values were calculated. Residues with smaller dRSA values corresponded to buried or conserved interfaces residues mediating stable interactions between spike chains. In contrast, those with higher dRSA values were more frequently found on the surface or at structurally dynamic interfaces between spike chains (Figure S6A). When dRSA values were compared with DMS expression scores, a qualitative agreement was apparent. This was most obvious within the NTD, where mutations in residues with lower dRSA values were likely to reduce protein expression, while mutations in residues with higher dRSA values were tolerated or even associated with increased expression (Figure S6B).

### NTD mutations compensate for RBD instability in Omicron variants

DMS data indicated that mutations altering spike protein expression dominantly contributed to viral infectivity. Previously we and others reported that immune evasion properties of Omicron were achieved mainly by RBD mutations while NTD mutations had more modest impact (Cui et al., 2022; Ikemura *et al*., 2022). Next, we examined the contribution of Omicron NTD mutations on infectivity and spike protein expression. We generated chimera spikes (C1-C6) with Wuhan-Hu-1 (D614G) and Omicron (BA.1) strains (Figure 5A) and examined the infectivity and spike expression. In the chimera spike with Wuhan background, Omicron NTD replacement substantially increased infectivity, whereas Omicron RBD replacement reduced it. In the case of Omicron background, Wuhan NTD replacement rendered the virus less infective, but Wuhan RBD replacement enhanced infectivity (Figure 5B). The same trend was observed in the spike protein expression on the cell surface as well as pseudovirus particles, where Omicron NTD and Wuhan RBD increased the spike protein expression, while Wuhan NTD and Omicron RBD reduced it (Figures 5C and 5D). These results indicated that Omicron RBD lost structural stability in exchange for gained immune evasion, and that the Omicron NTD evolution compensated for the RBD instability. Consistent with this hypothesis, when we generated the RBD-Fc protein, the Omicron RBD was low yield and highly prone to aggregation (Figure S7). To evaluate the stability of spike proteins, alteration of pseudovirus infectivity on plastic surface were measured over time. As expected, Wuhan with Omicron NTD (C1) exhibited stable infectivity, while Omicron with Wuhan NTD (C4) rendered the virus more fragile (Figure 5E).

**Figure 5.**
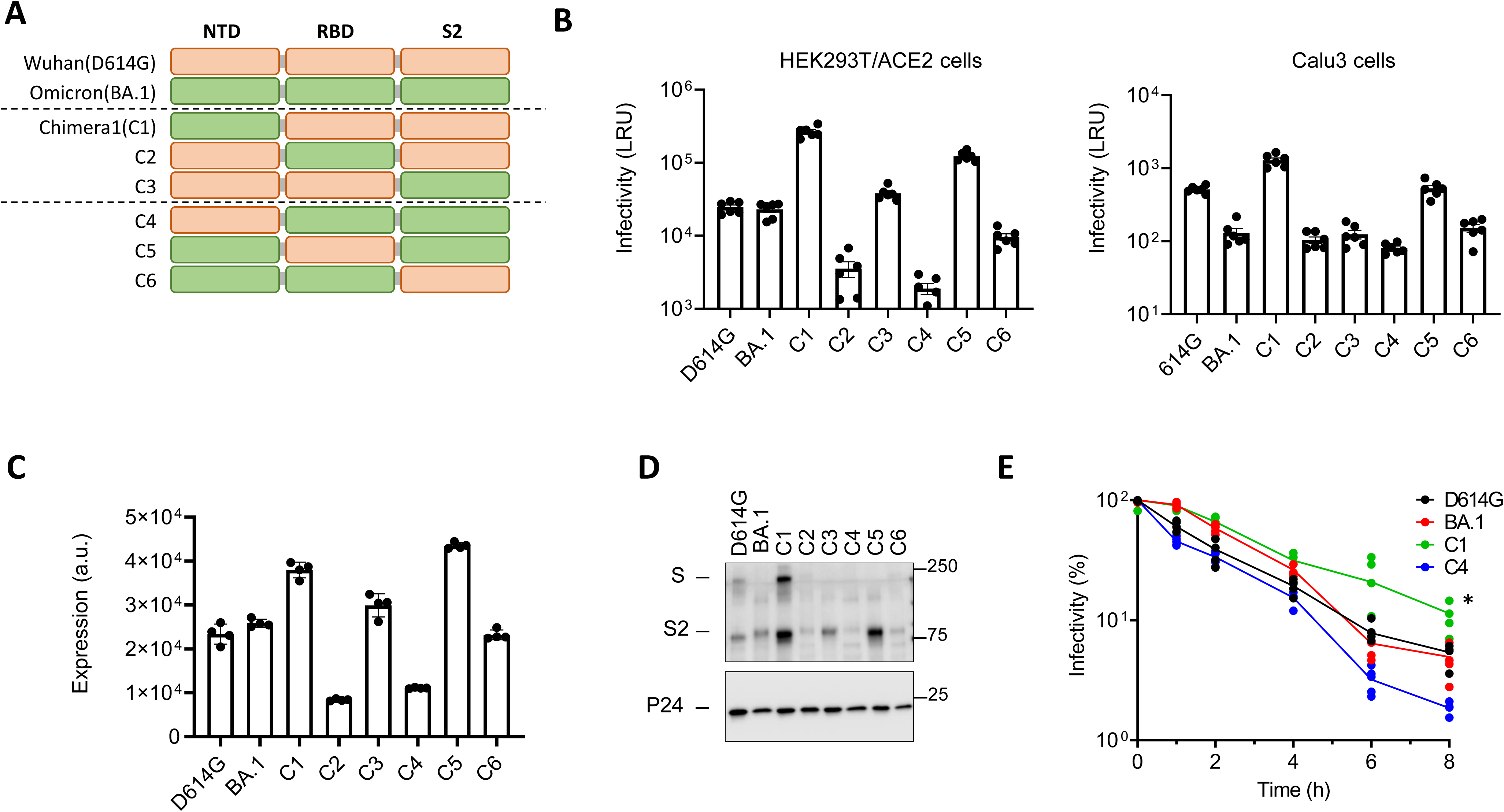
Omicron BA.1 NTD compensates the RBD structural instability. (A) Schematic of chimera spike used in this study. Spikes of Wuhan (D614G) and BA.1 were mixed in each domain and resultant chimera spikes were referred to as C1 to C6. (B) Infectivity of chimera spike-coating luciferase reporter pseudoviruses in HEK293T/ACE2 cells (left) and Calu-3 cells (right). (C) Spike expression of each chimera spike in HEK293T cells. Tagged HA was quantified by flow cytometry for anti HA antibody. (D) Spike protein in lentivirus particles pseudotyped with chimera spikes. The p24 lentivirus protein was used as an internal control. (E) Stability of pseudovirus particles on the surface of polystyrene plate. Virus infectivity was analyzed in each time point. * *P*=0.0004 vs. D614G, *P*=0.0002 vs.BA.1, *P*<0.0001 vs. C4. *P*-values were determined by one-way ANOVA and Dunnett’s multiple comparison test. For all plots, mean ± SD from at least two biological replicates.

To further understand the impact of each mutation on the Omicron NTD, we separated NTD mutations into 4 groups: NTD1, A67V/Δ69-70; NTD2, T95I; NTD3, G142D/Δ143-145; NTD4, Δ 211/L212I/ins214EPE (Figure 6A). When NTD group mutations were extracted from Omicron BA.1, NTD1, 2, and 4 exhibited impaired infectivity, but NTD3 did not (Figure 6B). In the case of introducing BA.1 NTD mutations in Wuhan-Hu-1 (D614G), NTD1,2, and 4 enhanced infectivity and NTD3 reduced it (Figure 6C). To further validate these observations, we also analyzed them in the chimeric Omicron spike with Wuhan NTD (C4) and confirmed no contribution of NTD3 to the enhanced infectivity (Figure 6D). Regarding spike protein expression, removal of NTD1, 2, and 4 reduced the expression, while their introduction into Wuhan NTD enhanced it in both Wuhan and C4 chimeric spike expression (Figures 6E-G). We previously reported that BA.1 NTD also modestly contributed to the escape from the neutralization of vaccinated sera(Ikemura *et al*., 2022). In neutralization assays with chimeric spike pseudoviruses, only NTD3 was responsible for immune evasion (Figures 6H). These intensive chimeric spike experiments revealed the evolutionary significance of Omicron BA.1 NTD, where NTD1, 2, 4 mutations compensated the spike stability and the NTD3 mutation contributed to immune evasion.

**Figure 6.**
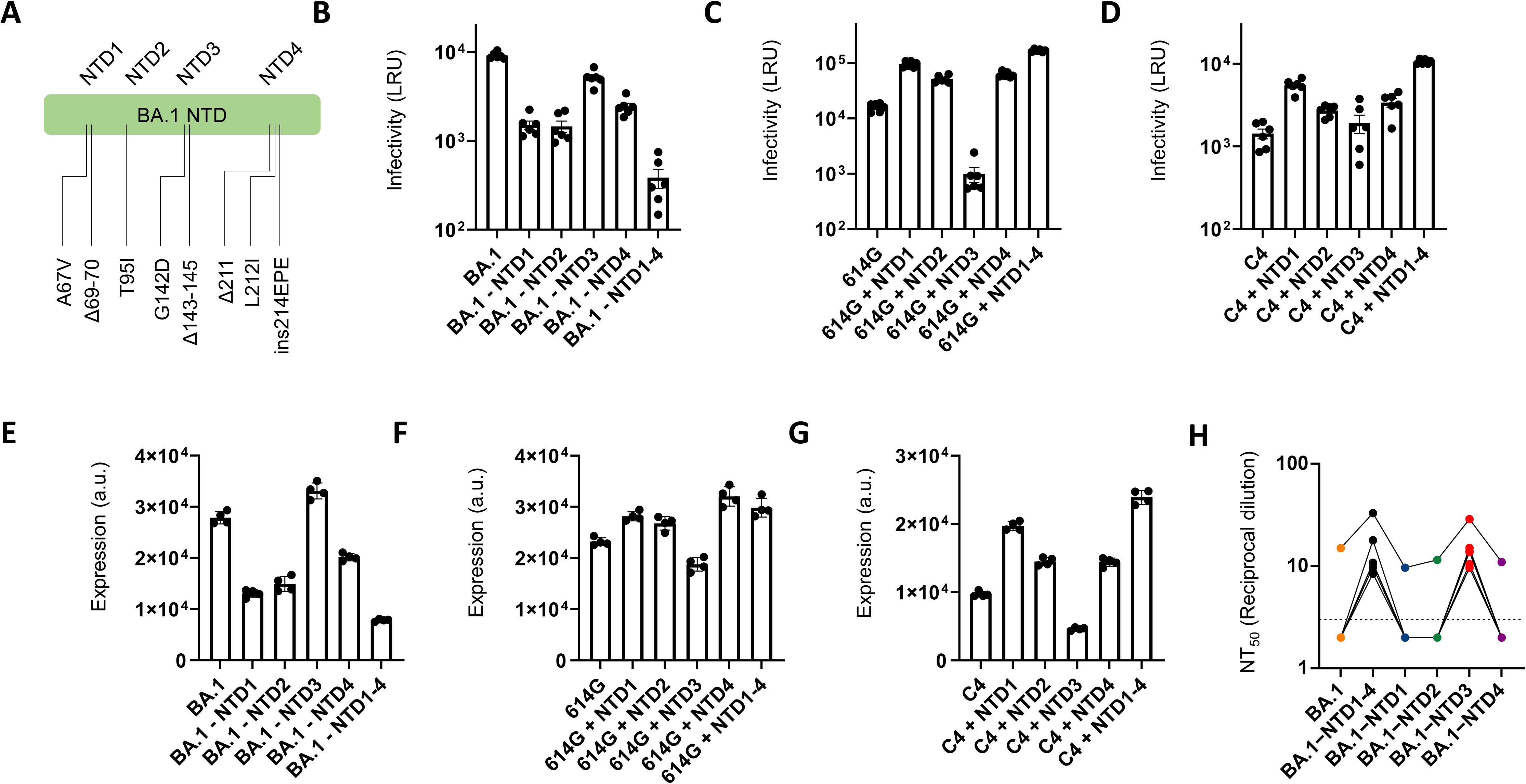
BA.1 NTD mutation enhances structural stability or immune evasion. (A) Schematic of BA.1 NTD mutation clusters used in this study. NTD1, A67V/Δ69-70; NTD2, T95I; NTD3, G142D/Δ143-145; NTD4, Δ211/L212I/ins214EPE. (B) Infectivity of luciferase reporter pseudoviruses carrying BA.1 spike with subtraction of NTD mutation cluster in HEK293T/ACE2 cells. (C) Infectivity of D614G variant spike with BA.1 NTD mutation cluster in HEK293T/ACE2 cells. (D) Infectivity of BA.1 chimeric spike with Wuhan NTD (C4) accompanied by BA.1 NTD mutation cluster in HEK293T/ACE2 cells. (E) Spike expression of BA.1 spike with subtraction of NTD mutation cluster in HEK293T cells. (F) Spike expression of D614G variant spike with BA.1 NTD mutation cluster in HEK293T cells. (G) Spike expression of BA.1 chimeric spike with Wuhan NTD (C4) accompanied by BA.1 NTD mutation cluster in HEK293T cells. For all plots above, mean ± SD from at least two biological replicates. (H) Neutralization efficacy in 293T/ACE2 cells for 5 vaccinated serum samples against BA.1 spike with subtraction of NTD mutation cluster. The NT50 values are summarized in the right panels. Dashed lines indicate detection limit.

We also conducted domain analysis in another Omicron variant, BA.2 with a similar lineup of chimeric spikes (Figure S8A). BA.2 NTD replacement enhanced both infectivity while spike expression and BA.2 RBD replacement conversely impaired them in Wuhan spike. Consistently, higher infectivity and spike expression were observed in BA.2 spike with Wuhan RBD, while BA.2 with Wuhan NTD exhibited the opposite phenotypes (Figures S8B and C). For further domain analysis, BA.2 NTD mutations were separated into 3 groups; NTD1, T19I/L24S/Δ25-27; NTD2, G142D; NTD3, V213G (Figure S8D). BA.2 NTD2 dominantly, and NTD3 modestly, improved the expression and infectivity, while NTD1 decreased both (Figures S8E-H). Interestingly, G142D with Δ143-145 enhanced immune evasion, but did not preserve infectivity, in BA.1; whereas, G142D alone strongly enhanced infectivity in BA.2.

### SARS-CoV-1 NTD contributes to higher infectivity

Although the spike RBD of SARS-CoV-2 has higher affinity to ACE2 than that of SARS-CoV-1(Lan et al., 2020; Nguyen et al., 2020; Shang et al., 2020), SARS-CoV-2 infectivity is much lower than SARS-CoV-1, at least in pseudovirus infection assays(Peacock et al., 2021; Zeng et al., 2020). SARS-CoV-2 is characterized by the gain of a furin cleavage site in the S1/S2 junction(Lan *et al*., 2020; Shang *et al*., 2020) and exhibits increased TMPRSS2-mediated infection and fusogenicity, which results in the promotion of virus transmissibility and pathogenicity(Peacock *et al*., 2021; Saito *et al*., 2022). In contrast, the biological meaning of alteration in the NTD remains poorly understood. To investigate this question, we constructed chimeric spikes of SARS-CoV1 and SARS-CoV-2 (Figure 7A). SARS-CoV-2 exhibited lower infectivity than SARS-CoV-1, consistent with other reports, and SARS-CoV-1 NTD replacement in SARS-CoV-2 improved infectivity. Conversely, SARS-CoV-1 infectivity was strikingly attenuated with SARS-CoV-2 NTD (Figure 7B). Spike expression was also altered in parallel with the infectivity (Figure 7C). Our binding affinity assay confirmed higher ACE2 affinity in the spike RBD of SARS-CoV-2 (Figure 7D). These results suggest that the NTD of SARS-CoV-1 was more fit to infect human cells.

**Figure 7.**
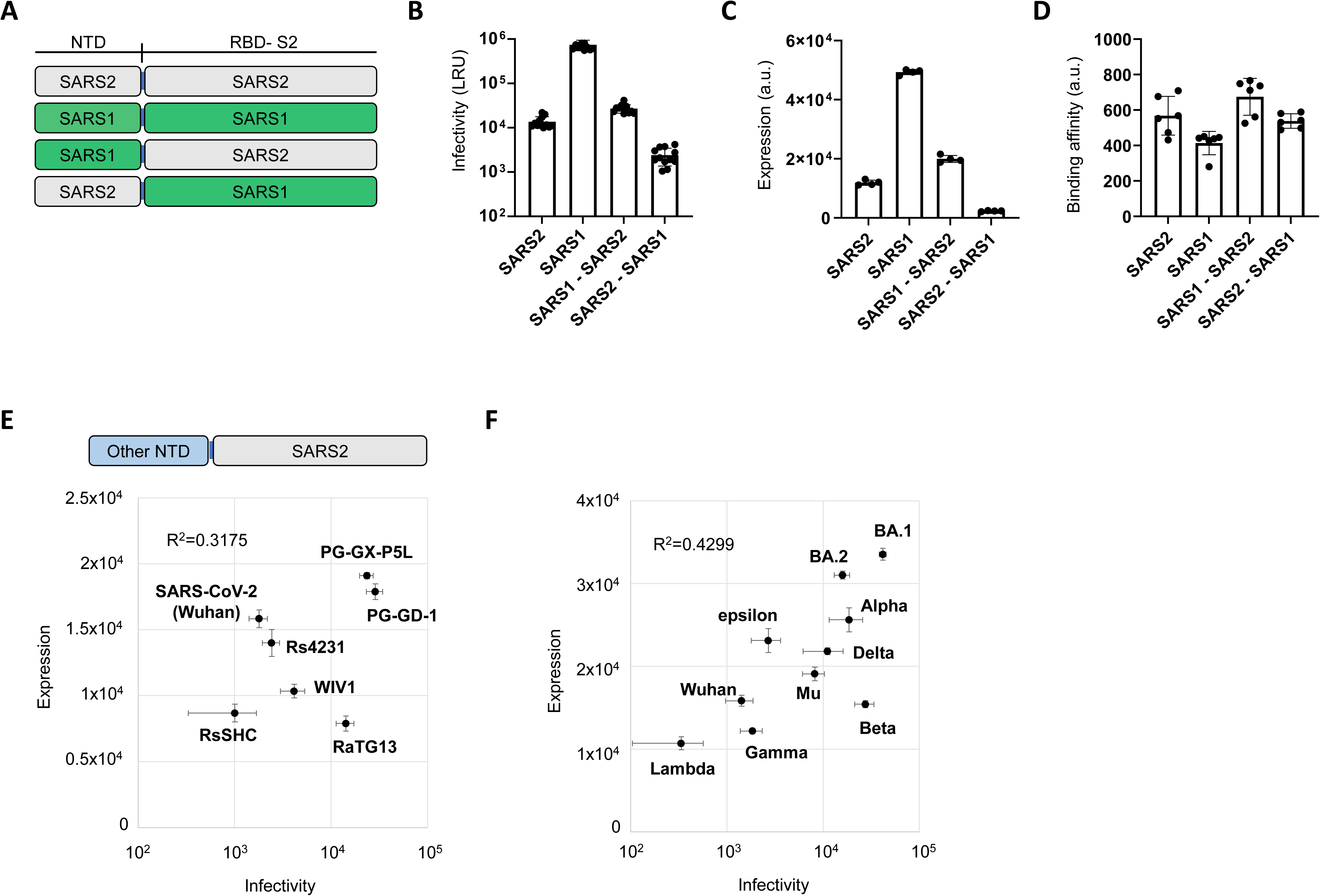
The fitness of NTDs from Sarbecoviruses and major SARS-CoV-2 variants. (A) Schematic of chimera spike with SARS-CoV-2 (Wuhan strain) and SARS-CoV-1. (B) Infectivity of each chimera spike-coating luciferase reporter pseudoviruses in HEK293T/ACE2 cells. (C) Spike expression of each chimera spike in HEK293T cells. (D) Spike affinity toward ACE2. Gating of flow cytometry-based affinity assessment was explained in Figure S1B. (E) Correlation between infectivity and spike expression in chimeric Wuhan D614G spike with Sarbecovirus NTDs. (F) Correlation between infectivity and spike expression in chimeric Wuhan D614G spike with major SARS-CoV-2 variant NTDs. The NTD sequences of sarbecoviruses and NTD mutations of SARS-CoV-2 variants are in Figure S9. For all plots, mean ± SD from at least two biological replicates.

Next, we examined the fitness of NTD from other sarbecoviruses and major SARS-CoV-2 variants in the context of chimera spike with Wuhan (Figures S9A-B). Correlation between spike expression and infectivity was also observed here and PG-GX-P5L and PG-GD-1 NTDs exhibited better fitness (Figure 7E). In the case of SARS-CoV-2 variants, BA.1 and BA.2 NTDs achieved the highest spike expression, indicating that the Omicron NTD further evolved to compensate the fragile RBD due to accumulated mutations for immune evasion (Figure 7F).

## Discussion

We have developed a DMS platform with an inverted infection assay to comprehensively evaluate viral infectivity, spike expression, affinity toward receptor and their correlation. The inverted infection assay uniquely employs ACE2-coating pseudovirus and spike-expressing cells. Naturally, SARS-CoV-2 spikes bind to cell surface ACE2 and protease-mediated cleavage of spike facilitates the membrane fusion between virus and cell. This process can be reproduced in the context of inverted positioning of spike and ACE2, and the expression of spike in cells allows to conduct high-throughput pooled library screening. Recently, pseudovirus-based spike DMS was performed to analyze infectivity and antibody escape but pseudotyped lentivirus DMS library was limited due to uncontrolled combination of mutations due to lentiviral swapping characteristics (Dadonaite *et al*., 2023). In comparison, our DMS assay system is all-encompassing and infectivity measurements similarly correlated with the enrichment of mutations in actual SARS-CoV-2 sequences. DMS analysis for SARS-CoV-2 has been intensively conducted by yeast surface display of the RBD and generated much data for RBD expression and binding to ACE2 or antibodies(Greaney *et al*., 2021; Starr *et al*., 2021a; Starr *et al*., 2021b; Starr *et al*., 2020). Although these DMS experiments provided important insights into the effect of viral mutations on infectivity and immune evasion, the region of study was limited to the RBD. In the context of these DMS data, the present study provides breadth and also reveals that spike protein expression is the primarily driving force for viral infectivity.

Based on DMS in the NTD, mutations enhancing spike expression and infectivity clustered in the loop regions and structurally fragile mutations were located at the interface to the RBD and S2. It is known that N1, N3 and N5 loop regions form an antigenic supersite and are targeted by neutralizing antibodies (Cerutti *et al*., 2021; McCallum *et al*., 2021). As the mutation frequency of naturally circulating mutants is high in the loop regions, it was expected that these mutations were selected by avoidance of neutralization by antibodies. However, they can also surprisingly increase spike expression and thus infectivity. Actually, in the case of BA.1 NTD, A67V/Δ69-70, T95I, and Δ211/L212I/ins214EPE mutations increase infectivity while G142D/Δ143-145 mutation contributes to immune evasion. A recent study also performed DMS on the NTD for spike protein expression in the context of full-length spike expression in HEK293T cells(Ouyang *et al*., 2022). Even though mutagenesis coverage was only 69%, this DMS similarly identified a trend wherein mutation tolerability was associated with distance to the RBD/S2. Omicron BA.1 contains several mutations in NTD loop regions and domain-wise experiments demonstrated that BA.1 NTD increased the environmental stability of the virus (Figure 5E). These results indicates that NTD loops are structurally flexible but, interestingly, a number of mutations facilitate increase in protein folding stability.

It is also surprising that spike protein expression is the predominant factor affecting viral infectivity rather than ACE2 affinity. Previously we generated high affinity ACE2 mutants to develop SARS-CoV-2 neutralizing ACE2 decoys(Higuchi *et al*., 2021; Ikemura *et al*., 2022). When this high affinity ACE2 was expressed in HEK 293T cells, pseudovirus infection rate was comparable to the wild-type ACE2 expression (Figure S3B). In another experiment, we isolated the mutant virus escaping wild-type ACE2 decoy and identified Y489H as the mutation responsible for this decoy escape. The Y489H mutation reduced ACE2 affinity and enabled escape from the ACE2 decoy. But this mutation also increased spike expression and overall infectivity was compensatively preserved(Urano et al., 2023). Similar decoy escape was reported in the case of CD4 decoys against HIV infection and it was suggested that nanoparticle coating CD4 could overcome the escape due to the avidity effect(Hoffmann et al., 2020). These results indicate the possibility that affinity between spike and ACE2 is saturated for effective infectivity and avidity effect due to higher spike expression dominantly determines viral infection activity.

The impact of NTD mutations on the spike expression and infectivity reinforces their contribution to SARS-CoV-2 evolution. The domain-wise analysis in Omicron BA.1 and BA.2 exhibited that Omicron RBD lost structural stability in exchange for immune evasion but the NTD evolved to preserve spike expression and infectivity. Interestingly, in the NTD of BA.1, only the G142D/Δ143-145 mutation exhibited no impact on the spike stability and simply contributed to antibody escape. BA.2 mutations in the N1 loop, T19I/L24S/Δ25-27, induced instability but mutations in this antigen supersite are also expected to affect immune evasion. Another study similarly reported domain-wise functional alteration in Omicron (Javanmardi et al., 2022), which made this evidence more convincing. In addition, we also examined the NTD of other major variants and found that BA.1, BA.2, and Alpha NTDs remarkably increased spike expression and infectivity. Moreover, NTD loop regions were poorly conserved among sarbecoviruses. When the NTD of SARS-CoV-2 was replaced with SARS-CoV-1, PG-GX-P5L or PG-GD-1, the infectivity was increased, indicating that SARS-CoV-2 could further evolve by recombination with zoonotic sarbecoviruses.

The DMS in the context of ACE2-coating pseudovirus infection into spike-expressing cells we describe here enables direct assessment of the alteration of infectivity due to single missense mutations. Combining with DMS analyzing spike expression and ACE2 affinity provides important insights into virus evolution, vaccine design and drug development. This system can also be applied to understand mutational escape from neutralizing reagents (Alcantara et al., 2023; Ikemura *et al*., 2022) and provides valuable information for antigenic surveillance.

## Acknowledgements

We would like to thank Takaaki Nakaya (Kyoto Prefectural University of Medicine) for helpful discussion; Kenzo Tokunaga (Department of Pathology, National Institute of Infectious Diseases) for the kind gift of the plasmid coding psPAX2-IN/HiBiT, This work was supported by the Japan Agency for Medical Research and Development (AMED), the Research Program on Emerging and Re-emerging Infectious Diseases under JP21fk0108465, JP21fk0108481, and 22fk0108523 (to A.H., J.T. and T.O.), the Platform Project for Supporting Drug Discovery and Life Science Research (Basis for Supporting Innovative Drug Discovery and Life Science Research) under JP21am0101075(2527) (to J.T.) and JP21am0101108 (to D.M.S), a grant from the Cell Science Research Foundation (to A.H.).

## Author contribution

A.H designed the research; S.T. and Y.H. conducted DMS experiments; S.M., Y.H. and N.I. performed pseudovirus neutralization assay; T.I. provided clinical serum samples from vaccinated individuals; S.L.,Y.K., D.M., and Y.O. performed and analyzed next-generation sequencing; T.A., J.T. purified and prepared the proteins; S.L. and D.M.S. manage SpikeScanDB; Y.I. and T.O. conducted authentic virus experiments; A.H., S.L. and D.M.S. wrote the manuscript; all authors discussed the results and commented on the manuscript.

## STAR Method

### Key resources table

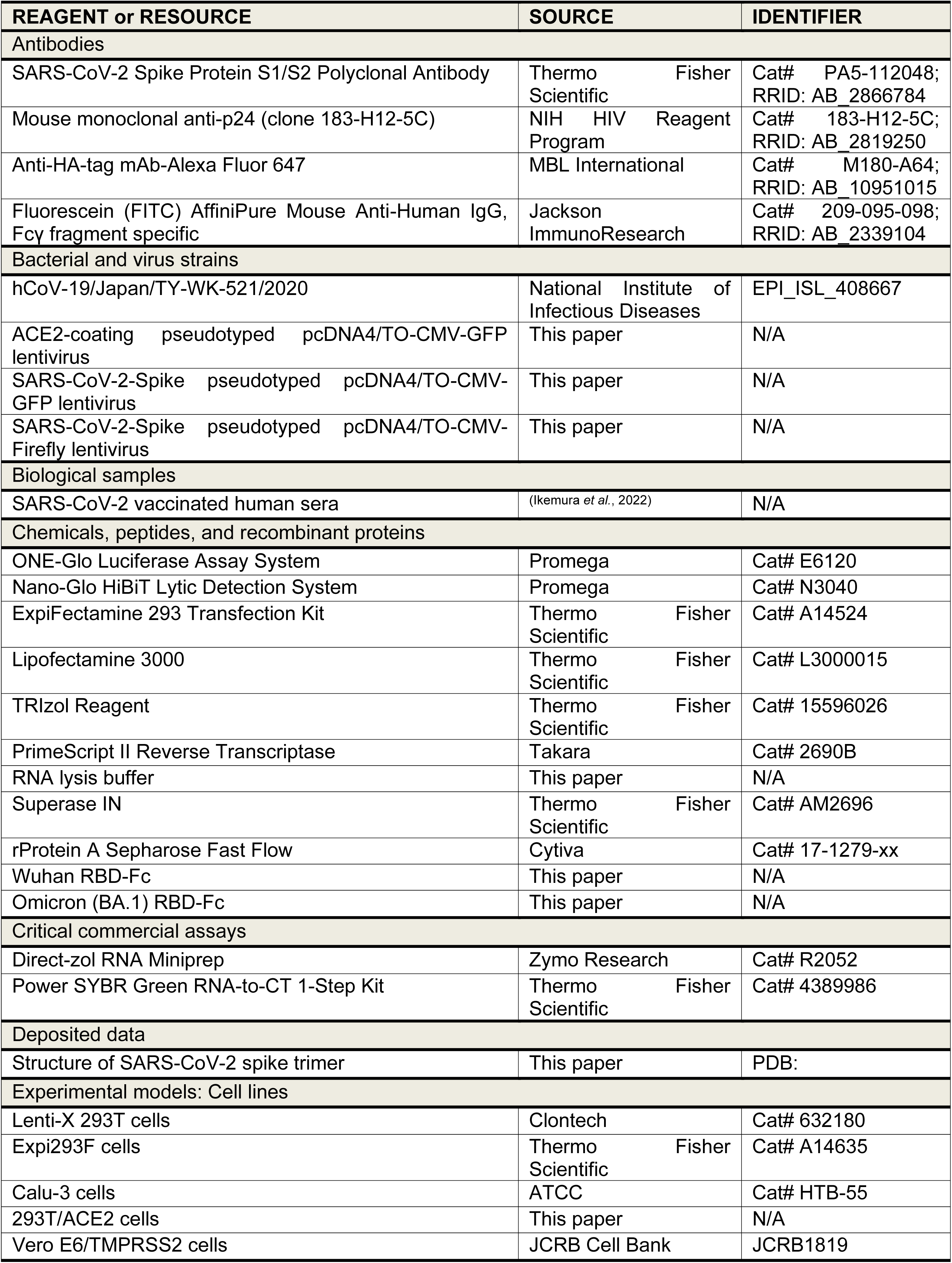
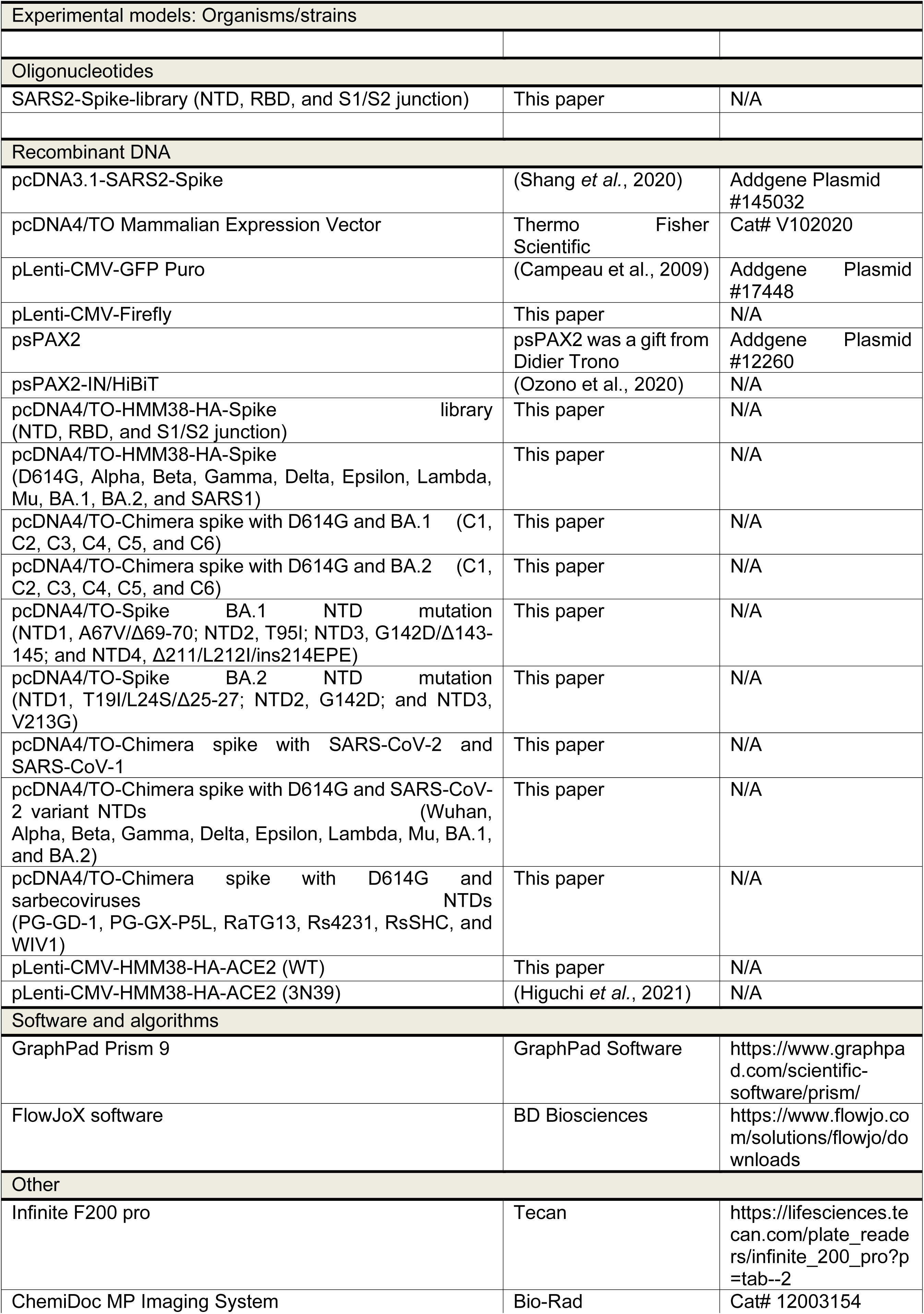
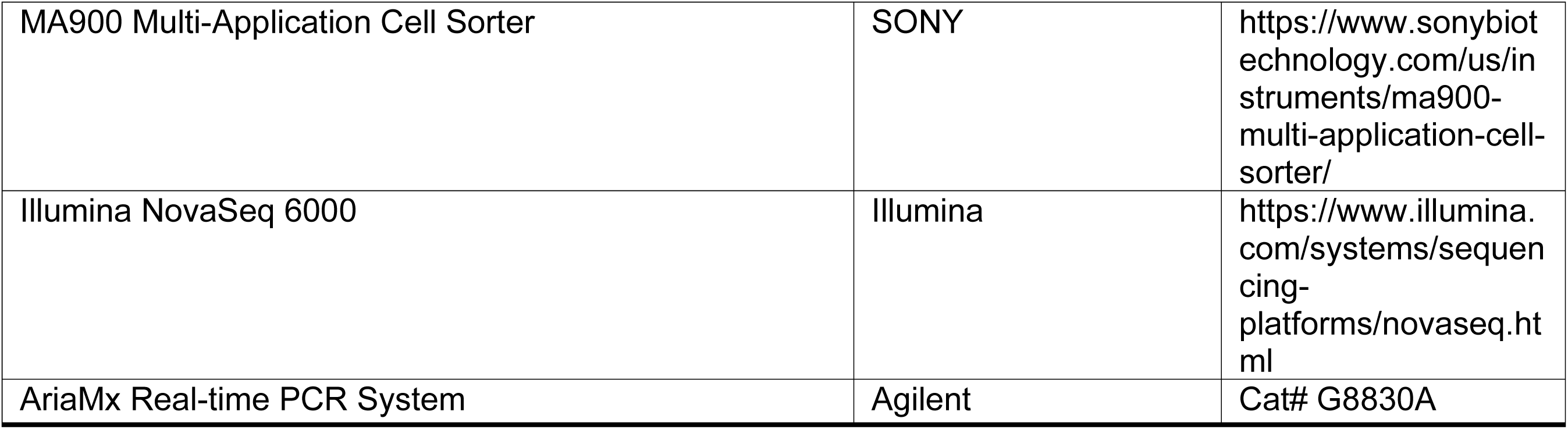

## RESOURCE AVAILABILITY

### Lead Contact

Further information and requests for resources and reagents should be directed to and will be fulfilled by the Lead Contact, Atsushi Hoshino.

### Material availability

All unique/stable reagents generated in this study are available from the Lead Contact with a completed Materials Transfer Agreement.

### Data and code availability

- Interactive versions of DMS heatmaps are at (https://sysimm.ifrec.osaka-u.ac.jp/sarscov2_dms/data/).
- All raw sequencing data are uploaded to the NCBI Short Read Archive under BioProject ID PRJNA988964
- This paper does not report original code.

## MATERIALS AND METHODS

### Study Design

Research blood samples were obtained from vaccinated individuals who were enrolled in a prospective cohort study approved by the Clinical Research Review Committee in Kyoto Prefectural University of medicine (ERB-C-1954-1). All human participants provided written informed consent. We collected peripheral blood from 7 individuals at 3 months post vaccination with two doses of the Pfizer-BioNTech vaccine, BNT162b2. The study size was determined by the number of samples that were available from the cohort study and not based on any power calculations. Experiments described in this manuscript were not performed blinded.

### Cell culture

Lenti-X 293T cells (Clontech) and its derivative, 293T/ACE2 cells were cultured at 37 °C with 5% CO_2_ in Dulbecco’s modified Eagle’s medium (DMEM, WAKO) containing 10% fetal bovine serum (Gibco) and penicillin/streptomycin (100 U/ml, Invitrogen). Calu3 cells were purchased from the American Tissue Culture Collection (ATCC, Rockville, MD) and cultured at 37 °C with 5% CO2 in DMEM/F-12 (WAKO) containing 10% fetal bovine serum (Gibco) and penicillin/streptomycin (100 U/ml, Invitrogen). All cell lines routinely tested negative for mycoplasma contamination.

### Plasmids construction

Using a plasmid encoding the SARS-CoV-2 S (addgene #145032) as a template, variant S fragments were amplified by bridge PCR with mutation-containing primers and cloned into pcDNA4TO (Invitrogen). To express HA-tagged spike protein, variant S fragments were cloned into pcDNA4TO as a form of HMM38-HA-spike. The NTD of Pangolin CoV GD-1, GX-P5L, Bat CoV RsSHC014, WIV1, RaTG13, and Rs4231 were synthesized (Integrated DNA Technologies) and cloned into pcDNA4TO (Invitrogen) in the form of ΔC19(10, 39). Pangolin CoV GX-P5L, Bat CoV RaTG13, and Rs4231 RBDs were synthesized (Integrated DNA Technologies) and cloned into the NTD of HMM38-HA-spike.

### Pseudotyped virus preparation and cell entry assays

Pseudotyped reporter virus assays were conducted as previously described (Higuchi *et al*., 2021). Spike protein-expressing pseudoviruses with a luciferase reporter gene was prepared by transfecting plasmids (pcDNA4TO spike, psPAX2-IN/HiBiT (Ozono *et al*., 2020), and pLenti firefly) into LentiX-293T cells with Lipofectamine 3000 (Invitrogen). After 48 hours, supernatants were harvested, filtered with a 0.45 μm low protein-binding filter (SFCA), and frozen at –80 °C. The lentivirus levels in viral supernatants were measured by the HiBiT assay, as previously described (Ozono *et al*., 2020). Briefly, a lentivirus stock with known levels of p24 antigen was serially diluted. Either the standards or viral supernatants containing pseudotyped viruses (25 μl) and LgBiT Protein (1:100)/HiBiT Lytic Substrate (1:50) in Nano-Glo HiBiT Lytic Buffer (25 μl) (Nano-Glo HiBiT Lytic Detection System; Promega) were mixed and incubated for 10min at room temperature according to the manufacturer’s instructions. HiBiT-based luciferase activity in viral supernatants was determined with Infinite F200 pro system (Tecan), and translated into p24 antigen levels. Pseudoviruses with different spike proteins corresponding to 1 ng of p24 antigen were incubated in 293T/ACE2 cells or Calu-3 cell. After 48 h, the cells were lysed in 100 μl of One-Glo Luciferase Assay Reagent (Promega). Firefly luciferase activity was determined with Infinite F200 pro system (Tecan).

### Pseudotyped virus neutralization assay

Pseudotyped reporter virus assays were conducted as previously described (Higuchi *et al*., 2021). Spike protein-expressing pseudoviruses with a luciferase reporter gene was prepared by transfecting plasmids (spikeΔC19 (Giroglou et al., 2004), psPAX2-IN/HiBiT (Ozono *et al*., 2020), and pLenti firefly) into LentiX-293T cells with Lipofectamine 3000 (Invitrogen). After 48 hours, supernatants were harvested, filtered with a 0.45 μm low protein-binding filter (SFCA), and frozen at –80 °C. The 293T/ACE2 cells were seeded at 10,000 cells per well in 96-well plates. HiBiT value-matched pseudoviruses and three-fold dilution series of serum were incubated for 1 hour, then this mixture was added to 293T/ACE2 cells. After 1 hour pre-incubation, medium was changed. At 48 hours post infection, cellular expression of the luciferase reporter, indicating viral infection, was determined using ONE-Glo Luciferase Assay System (Promega). Luminescence was read on Infinite F200 pro system (Tecan). The assay for each serum sample was performed in triplicate, and the 50% neutralization titer was calculated using Prism version 9 (GraphPad Software).

### Immunoblot

Pseudoviruses with different spike proteins corresponding to 1 ng of p24 antigen were loaded onto Tris-glycine sodium dodecyl sulfate-polyacrylamide gels, separated by electrophoresis, and then the proteins were transferred onto polyvinylidene difluoride membranes (Millipore, IPVH00010). The membranes were probed with an anti-SARS-CoV-2 spike protein S1/S2 polyclonal antibody (1:1,000, Thermo Fisher Scientific PA5-112048), and an anti-p24 monoclonal antibody (1:1,000). Blots were visualized with an HRP-conjugated secondary antibody, the Clarity Western ECL substrate (Bio-Rad Laboratories,1705060), and analyzed by ChemiDoc Touch MP (Bio-Rad).

### Protein synthesis and purification

Monoclonal antibodies and engineered ACE2 were expressed using the Expi293F cell expression system (Thermo Fisher Scientific) according to the manufacturer’s protocol. Fc-fusion proteins were purified from conditioned media using the rProtein A Sepharose Fast Flow (Cytiva). Fractions containing target proteins were pooled and dialyzed against phosphate buffered saline (PBS).

### Library construction, FACS, and Illumina sequencing analysis

Saturation mutagenesis was conducted independently in the NTD (S13 to T307), RBD (F329 to C538), and S1/S2 junction (F541 to S750) of ancestral original Wuhan strain. Pooled oligos with degenerate NNK codons were synthesized by Integrated DNA Technologies, Inc. Synthesized oligos were extended by overlap PCR and cloned into pcDNA4TO HMM38-HA-full length spike plasmids. Transient transfection conditions were used that typically provide no more than a single coding variant per cell(Chan *et al*., 2021). Expi293F cells at 2 × 10^6^ cells per ml were transfected with a mixture of 1 ng of library plasmid with 1 μg of pMSCV as a junk plasmid per ml using ExpiFectamine (Thermo Fisher Scientific). At 24 hours after transfection, cells were washed twice with PBS containing 10% bovine serum albumin (BSA) and then stained for 20 minutes with anti-hemagglutinin (HA) Alexa Fluor 647 (clone TANA2,1:4000 dilution; MBL) at 4 °C for expression assay. For affinity assay, cells were washed twice with PBS containing 10% BSA and then incubated with ACE2-Fc for stained for 20 minutes at 4 °C. Cells were again washed twice with PBS containing 10% BSA and then co-stained for 20 minutes with anti-HA Alexa Fluor 647 (clone TANA2,1:4000 dilution; MBL) and anti-human IgG-Fc γ fragment specific Fluorescein (FITC) (209-095-098, 1:100, Jackson ImmunoResarch) for 20 minutes at 4 °C. For infection assay, cells were incubated with ACE2-coating green fluorescent protein (GFP) reporter viruses, which were generated by transfecting pcDNA4TO ACE2, psPAX2 (addgene #12260), and pLenti GFP into LentiX-293T cells with Lipofectamine 3000 (Thermo Fisher Scientific). The viruses have ACE2 on the surface instead of glycoprotein and can infect the spike protein-expressing cells. At 24 hours after infection, cells were washed twice with PBS containing 10% BSA and then co-stained for 20 minutes with anti-HA Alexa Fluor 647 (clone TANA2,1:4000 dilution; MBL). Cells were again washed twice before sorting on a MA900 cell sorter (Sony). Dead cells, doublets, and debris were excluded by first gating on the main population by forward and side scatter. For expression analysis, the highest and lowest 25% HA signal among HA-positive cells were sorted. For affinity analysis, the highest and lowest 25% of cells with ACE2-Fc binding relative to HA-spike expression were sorted. For infection analysis, GFP-positive and -negative cells were collected from the HA positive (Alexa Fluor 647 positive) population. The total numbers of collected cells were about 2 million cells for each group. Total RNA was extracted from collected cells TRIzol (Thermo Fisher Scientific) and Direct-zol RNA MiniPrep (Zymo Research Corporation) according to the manufacturer’s protocol. First-strand complementary DNA (cDNA) was synthesized with PrimeScript II Reverse Transcriptase (Takara) primed with a gene-specific oligonucleotide. Libraries were designed for 3 sections separately in the RBD and then pooled. After cDNA synthesis, each library was amplified with specific primers. Following a second round of PCR, primers added adapters for annealing to the Illumina flow cell and sequencing primers, together with barcodes for experiment identification. The PCR products were sequenced on an Illumina NovaSeq 6000 using a 2 × 150 nucleotide paired-end protocol in Department of Infection Metagenomics, Research Institute for Microbial Diseases, Osaka University. Data were analyzed comparing the read counts with each group normalized relative to the wild type sequence read-count. Log10 enrichment ratios for all the individual mutations were calculated and normalized by subtracting the log10 enrichment ratio for the wild type sequence across the same PCR-amplified fragment.

### Viruses

The SARS-CoV-2 (Wuhan: 2019-nCoV/Japan/TY/WK-521/2020) strain was isolated at National Institute of Infectious Diseases (NIID). SARS-CoV-2 viruses were propagated in Vero E6/TMPRSS2 cells. Viral supernatant was harvested at two days post-infection and viral titers were determined by plaque assay.

### Quantitative RT-PCR of Viral RNA in the supernatant

The amount of RNA copies in the culture medium was determined using a qRT-PCR assay as previously described with slight modifications (Shema Mugisha et al., 2020). In brief, 5 µl of culture supernatants were mixed with 5 µl of 2× RNA lysis buffer (2% Triton X-100, 50 mM KCl, 100 mM Tris-HCl [pH 7.4], 40% glycerol, 0.4 U/µl of Superase IN [Thermo Fisher Scientific]) and incubated at room temperature for 10 minutes, followed by addition of 90 µl of RNase free water. Next, 2.5 µl of volume of the diluted samples was added to 17.5 µl of the reaction mixture. Real-time RT-PCR was performed with the Power SYBR Green RNA-to-CT 1-Step Kit (Applied Biosystems) using a AriaMx Real-Time PCR system (Agilent).

### Evaluation of virus stability on plastic surface

Virus survival was evaluated on polystyrene plate as previously described(Hirose et al., 2022). Luciferase reporter pseudoviruses with different spike proteins in 10 μL medium were applied on the surface of polystyrene plate. Each sample was incubated in a controlled environment (25°C, 45–55% relative humidity) for 0–8 h. The virus that remained on the surface was subsequently collected in 1.0 mL of DMEM and titrated by incubation in 293T/ACE2 cells.

### Statistical analysis

Neutralization measurements were done in technical triplicates and relative luciferase units were converted to percent neutralization and plotted with a non-linear regression model to determine 50% neutralization titer (NT_50_) values using GraphPad Prism software (version 9.0.0). Comparisons between two groups were made with two-sided unpaired *t* test. More than two groups were compared by one-way ANOVA and Tukey’s multiple comparison tests.

