## Supplementa figures for "Deep mutational scanning of whole SARS-CoV-2 spike in an inverted infection system"

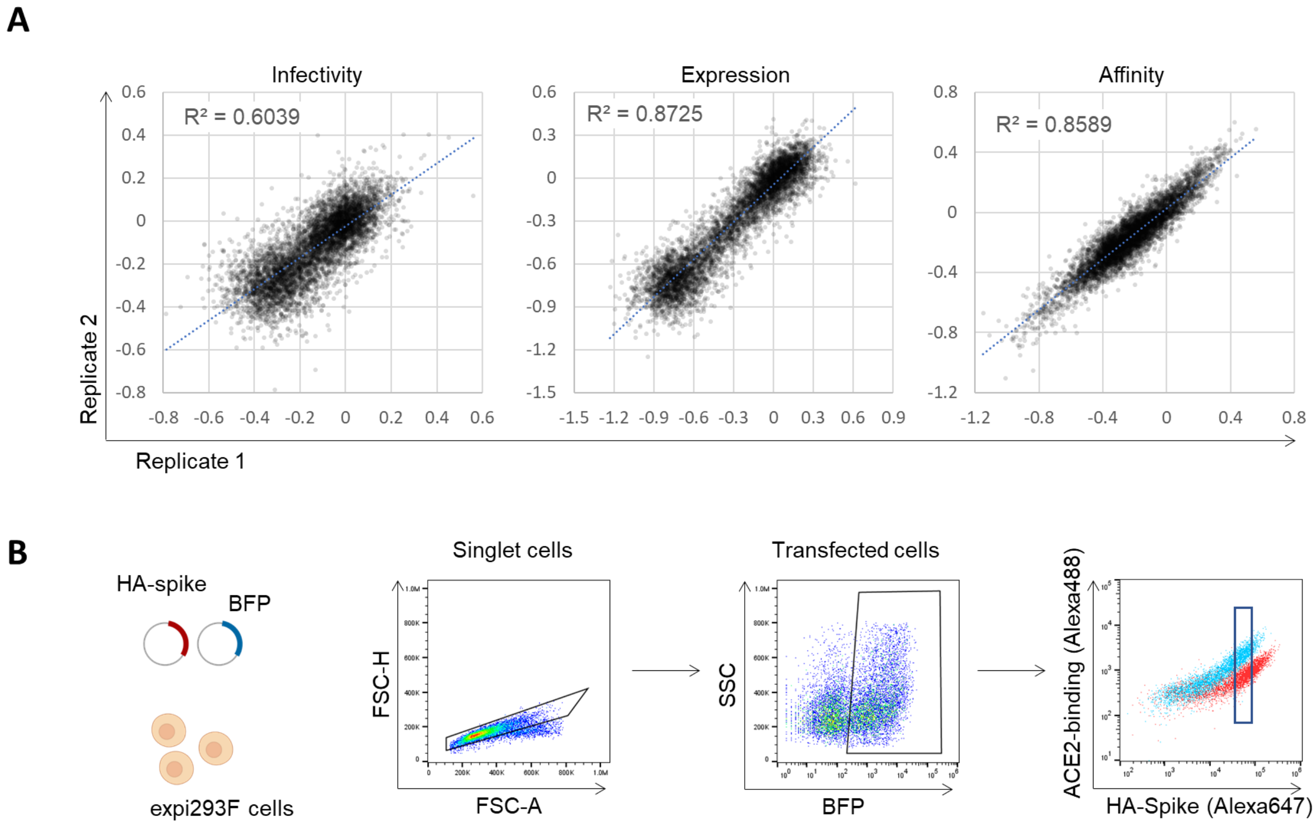


**Figure S1. Deep mutational scanning for the N-terminal domain.**

(A) Reproducibility of deep mutational scanning (DMS) for N-terminal domain (NTD) analyzing infectivity, spike expression and affinity toward ACE2 across replicates.

(B) Gating strategy to evaluate the ACE2 binding affinity in flow cytometry. The spike expressing cells were gated according to the co-transfected BFP signal. The signal of ACE2 bound to spike (Alexa 488) was quantified in arbitrary signal range of HA-spike expression (Alexa 647).

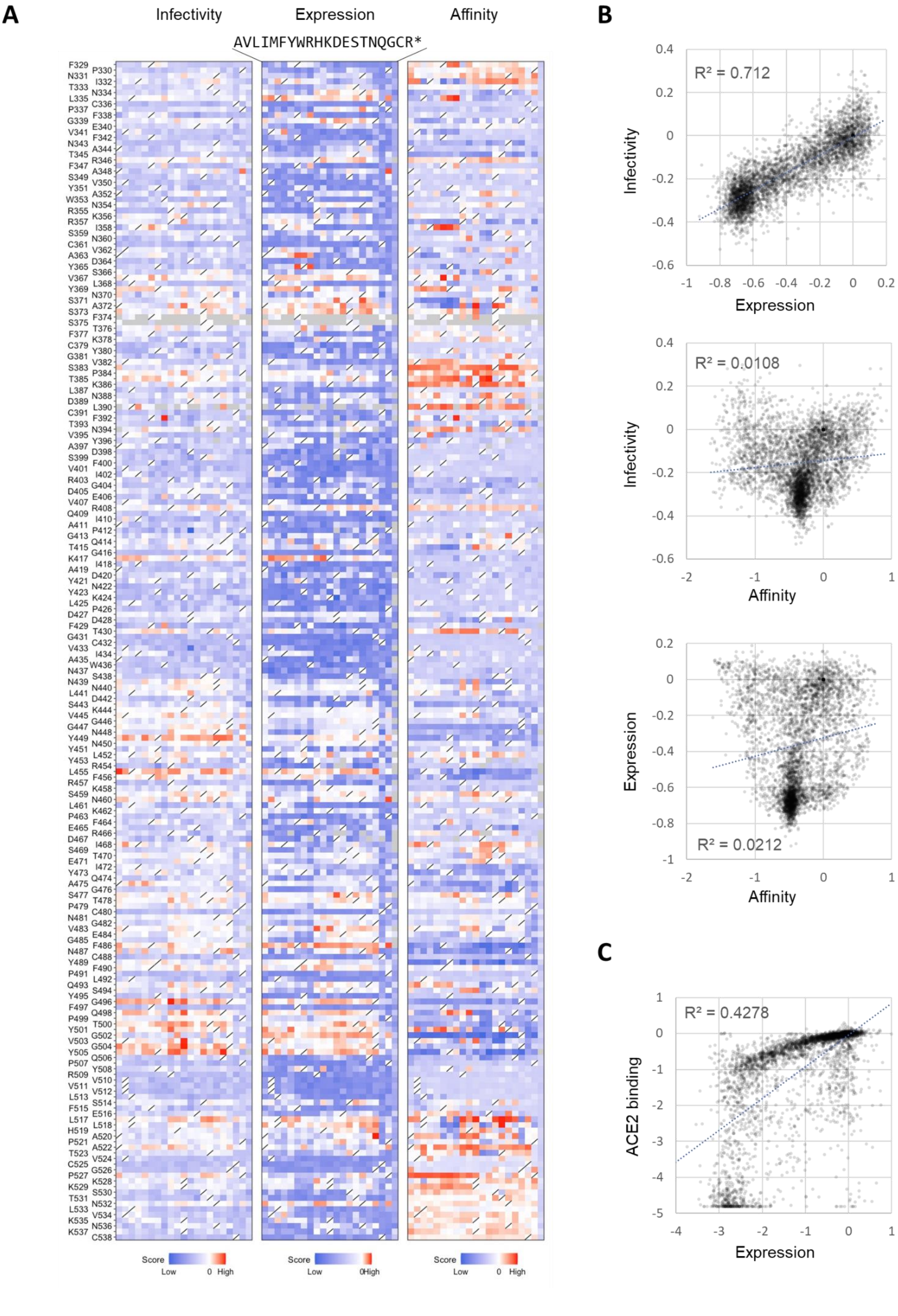
A
Infectivity
Expression
Affinity
AVLIMFYWRHKDESTNQGCR*
F329
P330
N331
I332
T333
N334
L335
C336
P337
F338
G339
E340
V341
F342
N343
A344
T345
R346
F347
A348
S349
V350
Y351
A352
W353
N354
R355
K356
R357
I358
S359
N360
C361
V362
A363
D364
Y365
S366
V367
L368
Y369
N370
S371
A372
S373
F374
S375
T376
F377
K378
C379
Y380
G381
V382
S383
P384
T385
K386
L387
N388
D389
L390
C391
F392
T393
N394
V395
Y396
A397
D398
S399
F400
V401
I402
R403
G404
D405
E406
V407
R408
Q409
I410
A411
P412
G413
Q414
T415
G416
K417
I418
A419
D420
Y421
N422
Y423
K424
L425
P426
D427
D428
F429
T430
G431
C432
V433
I434
A435
W436
N437
S438
N439
N440
L441
D442
S443
K444
V445
G446
G447
N448
Y449
N450
Y451
L452
Y453
R454
L455
F456
R457
K458
S459
N460
L461
K462
P463
F464
E465
R466
D467
I468
S469
T470
E471
I472
Y473
Q474
A475
G476
S477
T478
P479
C480
N481
G482
V483
E484
G485
F486
N487
C488
Y489
F490
P491
L492
Q493
S494
Y495
G496
F497
Q498
P499
T500
Y501
G502
V503
G504
Y505
Q506
P507
Y508
R509
V510
V511
V512
L513
S514
F515
E516
L517
L518
H519
A520
P521
A522
T523
V524
C525
G526
P527
K528
K529
S530
T531
N532
L533
V534
K535
N536
K537
C538
Score Low 0 High
Score Low 0High
Score Low 0 High
B
R² = 0.712
Infectivity
Expression
0.4
0.2
0
-0.2
-0.4
-0.6
-1
-0.8
-0.6
-0.4
-0.2
0
0.2
R² = 0.0108
Infectivity
Affinity
-2
-1
0
1
R² = 0.0212
Expression
Affinity
C
R² = 0.4278
ACE2 binding
Expression
1
0
-1
-2
-3
-4
-5

**Figure S2. Deep mutational scanning for the receptor binding domain.**

(A) Heatmap showing how all single mutations affect the infectivity, spike expression, and affinity toward ACE2 in the receptor binding domain (RBD). Squares are colored by mutational effect according to scale bars on the bottom, with blue indicating deleterious mutations. Squares with a diagonal line through them indicate the original Wuhan strain amino acid.

(B) Correlation between functional alterations of infectivity, spike expression and affinity from DMS in the RBD.

(C) Scatter plot showing the correlation between ACE2 binding and expression in Starr et al. DMS of yeast display-based RBD analysis (1).

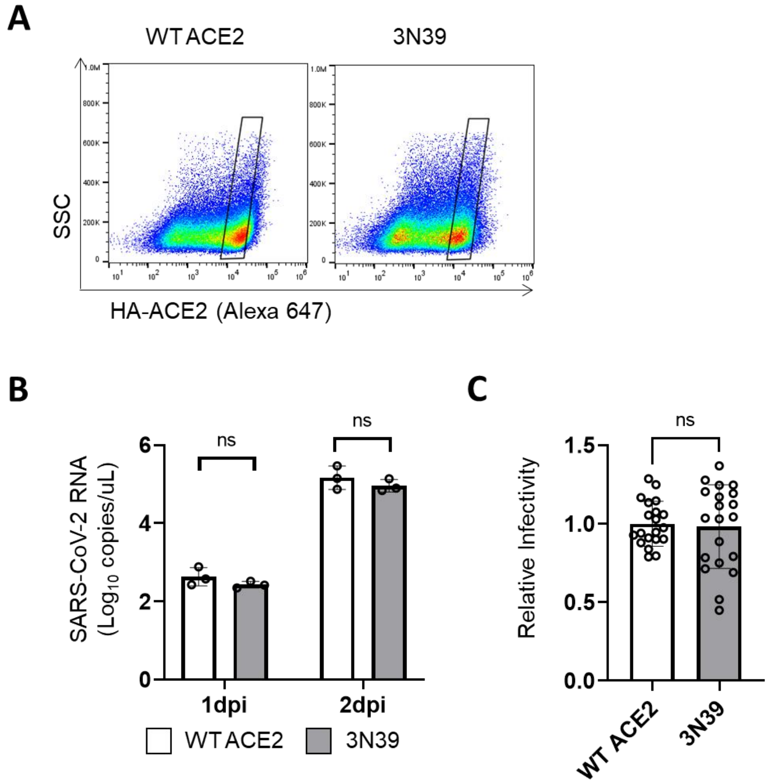


**Figure S3. Strongly increased affinity showed no enhancement of virus infectivity.**

(A) Gating of cell sorting to prepare cells similarly expressing ACE2 proteins.

(B) Comparison of viral growth between wild-type (WT) ACE2 and 3N39 expressing HEK293T cells at 1-day and 2-day post-infection. Viral RNA in supernatant was quantified by qPCR. Data are mean ± SD from three technical replicates.

(C) The infectivity of D614G pseudovirus in HEK293T cells expressing WT ACE2 and 3N39. Data are mean ± SD from three biological replicates.

*P*-values were determined by two-sided unpaired *t*-test.

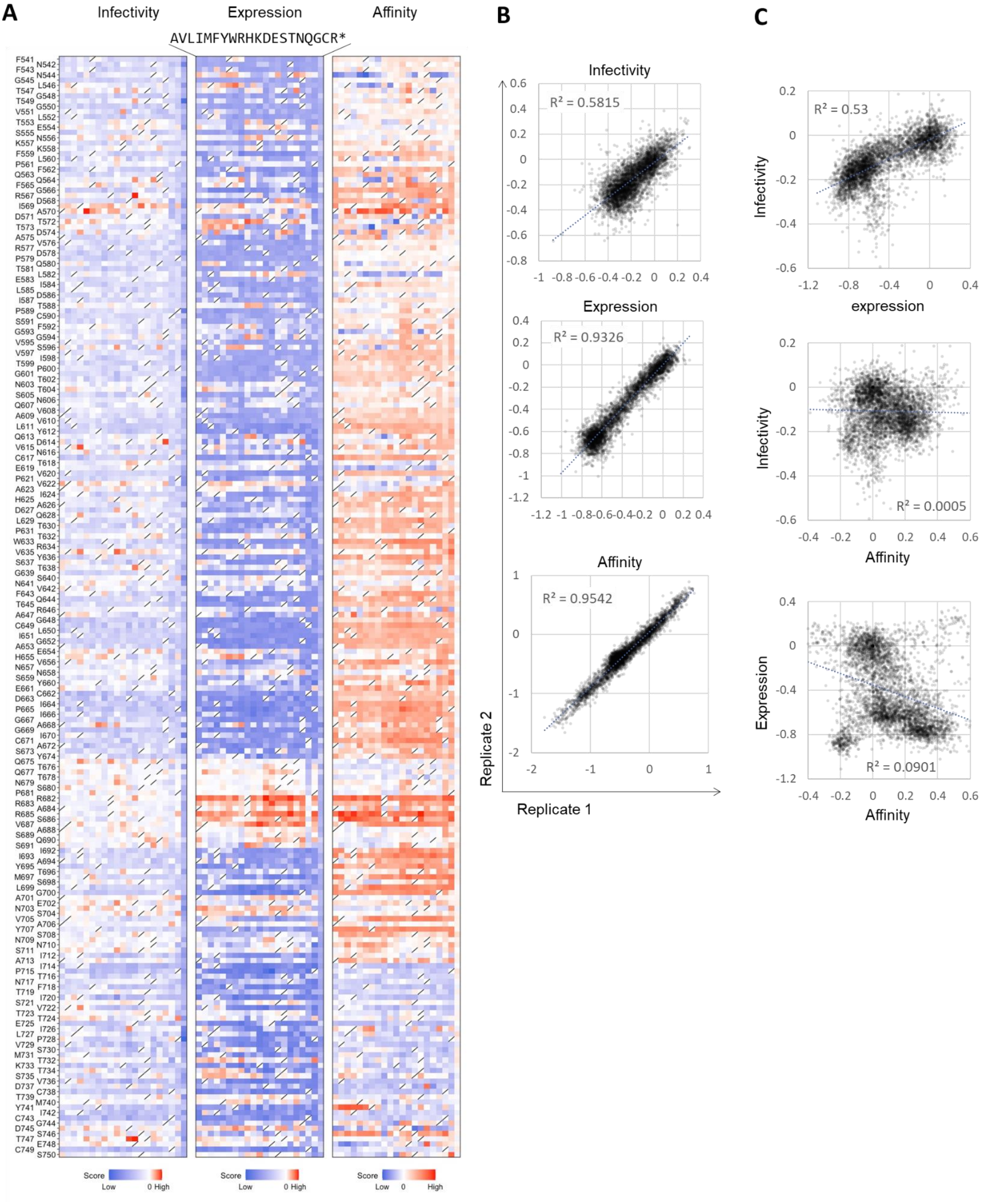
A
Infectivity
Expression
Affinity
AVLIMFYWRHKDESTNQGCR*
Score Low 0 High
Score Low 0 High
Score Low 0 High
B
Infectivity
R² = 0.5815
Expression
R² = 0.9326
Affinity
R² = 0.9542
Replicate 2
Replicate 1
C
R² = 0.53
Infectivity
expression
Infectivity
R² = 0.0005
Affinity
Expression
R² = 0.0901
Affinity

**Figure S4 Deep mutational scanning for the S1/S2 junction.**

(A) Heatmap showing how all single mutations affect the infectivity, spike expression, and affinity toward ACE2 in the S1/S2 junction region. Squares are colored by mutational effect according to scale bars on the bottom, with blue indicating deleterious mutations. Squares with a diagonal line through them indicate the original Wuhan strain amino acid.

(B) Correlation in mutation effects on the alteration of infectivity, spike expression and affinity toward ACE2 across replicates.

(C) Correlation between functional alterations of infectivity, spike expression and affinity.

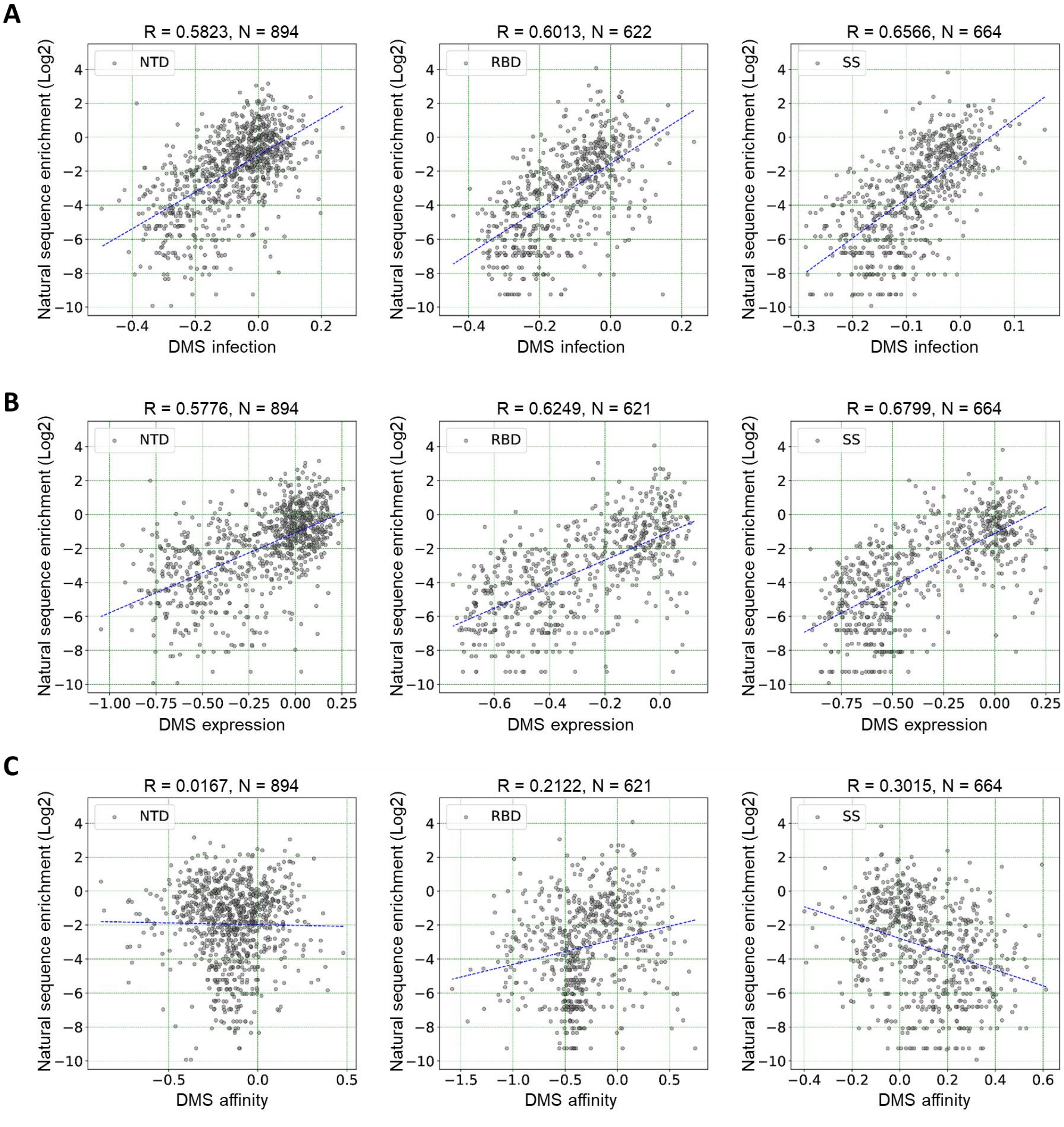


**Figure S5 Functional effects of each domain mutations on the viral transmission fitness**

(A-C) Correlation between the mutation frequency of circulating SARS-CoV-2 and functional alteration of spike including infectivity (A), spike expression (B), and affinity toward ACE2 (C) in the NTD, RBD, and S1/S2 junction.

A

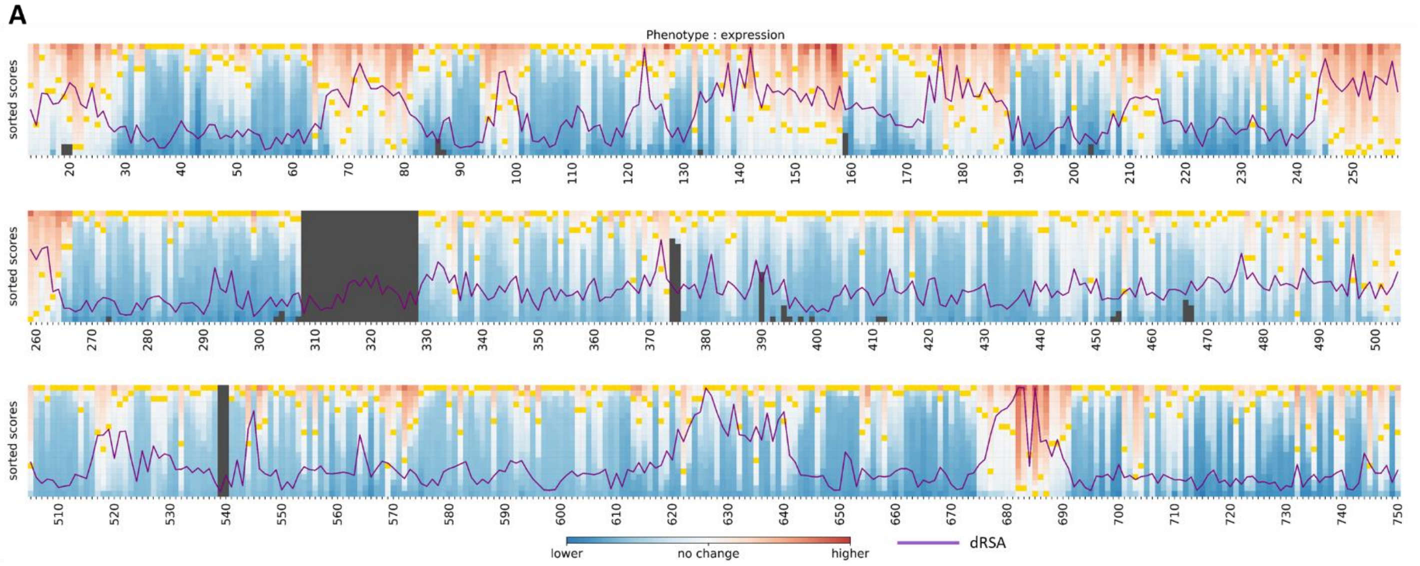


B

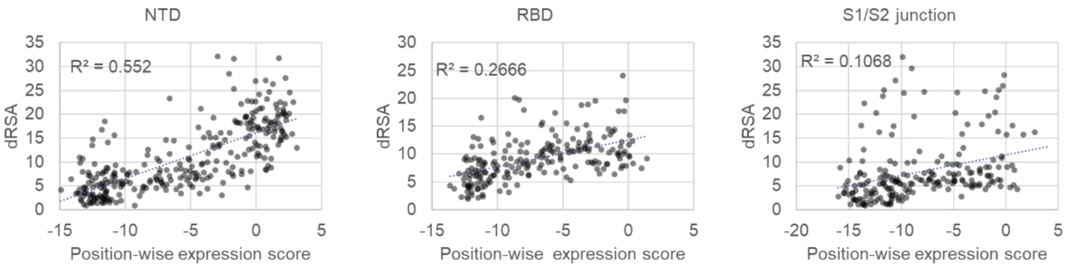


**Figure S6 Alteration of spike expression is associated with solvent accessibility**

(A) Heatmap showing spike expression and standard deviation of RSA (dRSA; purple line). Each single mutation value was sorted to visualize the position-wise score for the enhancement of spike expression.

(B) Correlation between position-wise spike expression score and dRSA in the NTD, RBD, and S1/S2 junction.

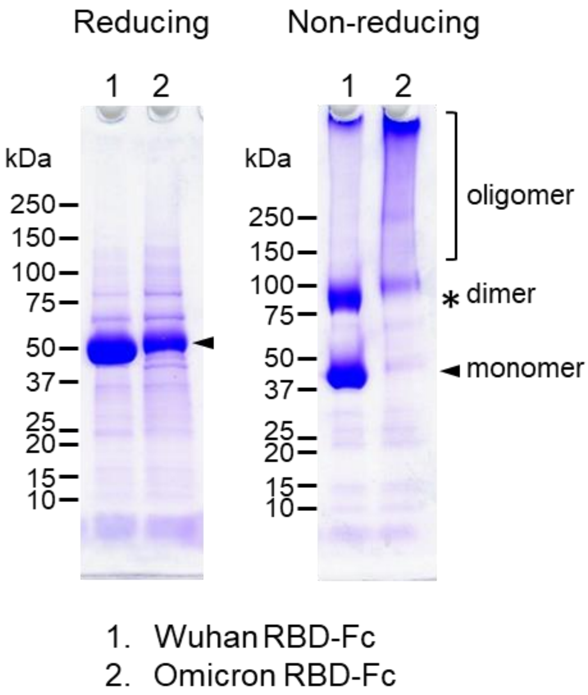


**Figure S7 Structural instability of Omicron RBD**

SDS-PAGE of purified RBD proteins. Omicron RBD exhibited high level of oligomer formation.

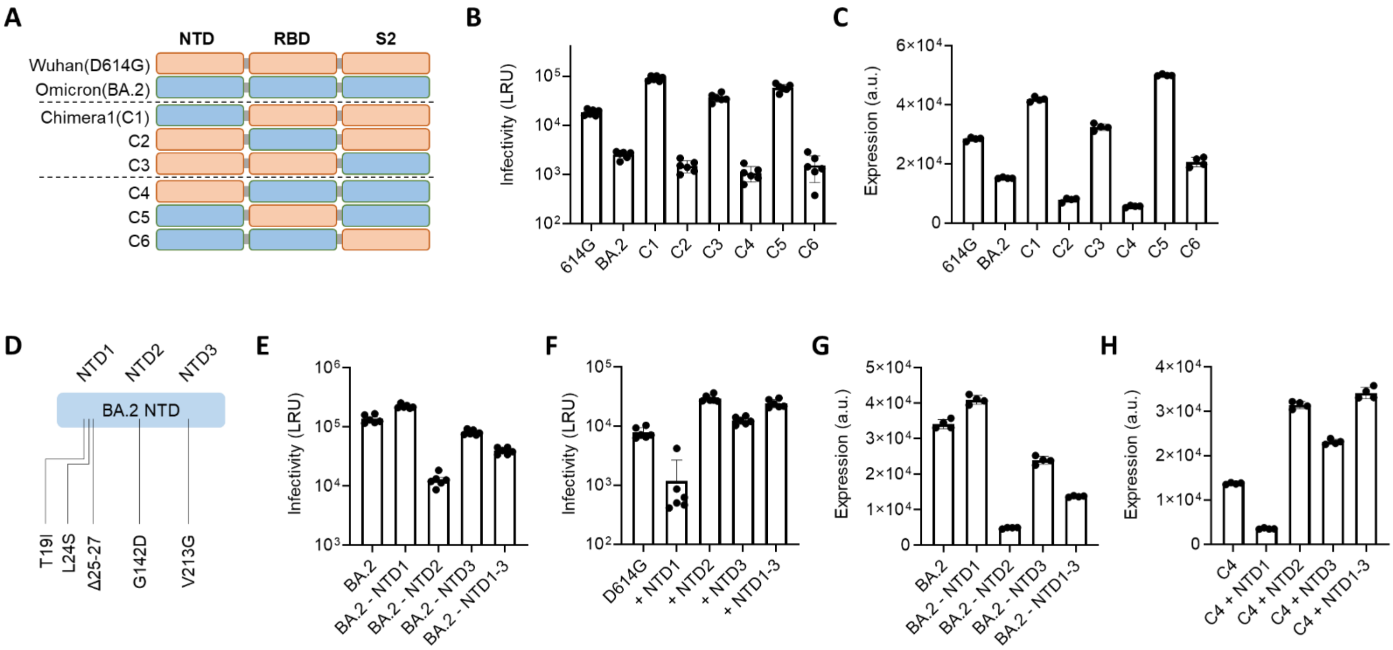


**Figure S8 BA.2 NTD mutation enhances structural stability.**

(A) Schematic of chimera spike used in this study. Wuhan and BA.2 were mixed in each domain.

(B) Infectivity of chimera spike-coating luciferase reporter pseudoviruses in HEK293T/ACE2 cells.

(C) Spike expression of each chimera spike in HEK293T cells.

(D) Schematic of BA.2 NTD mutation clusters used in this study. NTD1, T19I/L24S/Δ25-27; NTD2, G142D; NTD3, V213G.

(E) Infectivity of BA.2 spike with subtraction of NTD mutation cluster in HEK293T cells.

(F) Infectivity of D614G variant spike with BA.2 NTD mutation cluster in HEK293T cells.

(G) Spike expression of BA.2 spike with subtraction of NTD mutation cluster in HEK293T cells.

(H) Spike expression of D614G variant spike with BA.2 NTD mutation cluster in HEK293T cells.

For all plots, mean ± SEM from at least two biological replicates.

**A**

```
WIV1          YTIEKCLDFDDRTPPANTQFLSSHRGVYYYPDDIFRSNVLHLVQDHFLPFDSNVTRFITFG   60
RsSHC014      YTIEKCLDFDDRTPPANTQFLSSHRGVYYYPDDIFRSNVLHLVQDHFLPFDSNVTRFITFG   60
SARS-CoV      SDLDRCTTFDDVQAPNYTQHTSSMRGVYYYPDEIFRSDTLYLTQDLFLPFYSNVTGFHTIN   60
Rs4231        SDLESCTTFDDVQAPNYPQHSSSRRGVYYYPDEIFRSDTLYLTQDLFLPFYSNVTGFHTIN   60
PG-GD-1       ----QCVNLTGRAAIQPSFTNSSQRGVYYYPDTIFRSNTLVLSQGYFLPFYSNVSWYYALT   56
PG-GX-P5E     ----QCVNLTTRTGIPPGYTNSSTRGVYYYPDKVFRSSILHLTQDLFLPFFSNVTWFNTIT   56
Wuhan         ----QCVNLTTRTQLPPAYTNSFTRGVYYYPDKVFRSSVLHSTQDLFLPFFSNVTWFHAIH   56
RaTG13        ----QCVNLTTRTQLPPAYTNSSTRGVYYYPDKVFRSSVLHLTQDLFLPFFSNVTWFHAIH   56
                   *  :            *  ******* :***. *   *. **** ***: : ::

WIV1          -------LNFDNPIIPFKDGIYFAATEKSNVIRGWVFGSTMNNKSQSVIIMNNSTNLVIR   113
RsSHC014      -------LNFDNPIIPFRDGIYFAATEKSNVIRGWVFGSTMNNKSQSVIIMNNSTNLVIR   113
SARS-CoV      -------HTFDNPVIPFKDGIYFAATEKSNVVRGWVFGSTMNNKSQSVIIINNSTNVVIR   113
Rs4231        -------HRFDNPVIPFKDGVYFAATEKSNVVRGWVFGSTMNNKSQSVIIINNSTNVVIR   113
PG-GD-1       KTN-SAEKRVDNPVLDFKDGIYFAATEKSNIVRGWIFGTTLDNTSQSLLIVNNATNVIIK   115
PG-GX-P5E     YQG--GSKKFDNPVLPFNDGVYFASTEKSNIIRGWIFGTTLDARTQSLLIVNNATNVVIK   114
Wuhan         VSGTNGTKRFDNPVLPFNDGVYFASTEKSNIIRGWIFGTTLDSKTQSLLIVNNATNVVIK   116
RaTG13        VSGTNGIKRFDNPVLPFNDGVYFASTEKSNIIRGWIFGTTLDSKTQSLLIVNNATNVVIK   116
                       .***:: *.**:***:*****::***:**:*::  :**::*:**:**::*:

WIV1          ACNFELCDNPFFVVLKSNNTQI----PSYIFNNAFNCTFEYVSKDFNLDLGEKPGNFKDL   169
RsSHC014      ACNFELCDNPFFVVLKSNNTQI----PSYIFNNAFNCTFEYVSKDFNLDLGEKPGNFKDL   169
SARS-CoV      ACNFELCDNPFFAVSKPMGTQT----HTMIFDNAFNCTFEYISDAFSLDVSEKSGNFKHL   169
Rs4231        ACNFELCDNPFFAVSKPTGTQT----HTMIFDNAFNCTFEYISDSFSLDVAEKSGNFKHL   169
PG-GD-1       VCNFQFCYDPYLSGYYHN-NKTWSTREFAVYSSYANCTFEYVSKSFMLDIAGKSGLFDTL   174
PG-GX-P5E     VCEFQFCTDPFLGVYYHNNNKTWVENEFRVYSSANNCTFEYISQPFLMDLEGKQGNFKNL   174
Wuhan         VCEFQFCNDPFLGVYYHKNNKSWMESEFRVYSSANNCTFEYVSQPFLMDLEGKQGNFKNL   176
RaTG13        VCEFQFCNDPFLGVYYHKNNKSWMESEFRVYSSANNCTFEYVSQPFLMDLEGKQGNFKNL   176
              .*:*::* :*::       .:        ::..  ******:*. * :*:  * * *. *

WIV1          REFVFRNKDGFLHVYSGYQPISAASGLPTGFNALKPIFKLPLGINITNFRTLLTAFPP--   227
RsSHC014      REFVFRNKDGFLHVYSGYQPISAASGLPTGFNALKPIFKLPLGINITNFRTLLTAFPP--   227
SARS-CoV      REFVFKNKDGFLYVYKGYQPIDVVRDLPSGFNTLKPIFKLPLGINITNFRAILTAFLP--   227
Rs4231        REFVFKNKDGFLYVYKGYQPIDVVRDLPSGFNILKPIFKLPLGINITNFRAILTAFLP--   227
PG-GD-1       REFVFRNVDGYFKIYSKYTPVNVNSNLPIGFSALEPLVEIPAGINITKFRTLLTIHRGDP   234
PG-GX-P5E     REFVFKNVDGYFKIYSKHTPIDLVRDLPRGFAALEPLVDLPIGINITRFQTLLALHRSYL   234
Wuhan         REFVFKNIDGYFKIYSKHTPINLVRDLPQGFSALEPLVDLPIGINITRFQTLLALHRSYL   236
RaTG13        REFVFKNIDGYFKIYSKHTPINLVRDLPPGFSALEPLVDLPIGINITRFQTLLALHRSYL   236
              *****:* **:: :*. : *:.   .** **  *:*:..:* *****.*:::*: .

WIV1          ----RPDYWGTSAAAYFVGYLKPTTFMLKYDENGTITDAVDCSQNPLAELKCSVKS       279
RsSHC014      ----RPDYWGTSAAAYFVGYLKPTTFMLKYDENGTITDAVDCSQNPLAELKCSVKS       279
SARS-CoV      ----AQDTWGTSAAAYFVGYLKPTTFMLKYDENGTITDAVDCSQNPLAELKCSVKS       279
Rs4231        ----AQDTWGTSAAAYFVGYLKPATFMLKYDENGTITDAVDCSQNPLAELKCSVKS       279
PG-GD-1       MP---NNGWTVFSAAYYVGYLAPRTFMLNYNENGTITDAVDCALDPLSEAKCTLKS       287
PG-GX-P5E     TPGKLESGWTTGAAAYYVGYLQQRTFLLSYNQNGTITDAVDCSLDPLSETKCTLKS       290
Wuhan         TPGDSSSGWTAGAAAYYVGYLQPRTFLLKYNENGTITDAVDCALDPLSETKCTLKS       292
RaTG13        TPGDSSSGWTAGAAAYYVGYLQPRTFLLKYNENGTITDAVDCALDPLSETKCTLKS       292
                    . * . :***:****   **:*.*::**********: :**:* **::**
```

**B**

| Alpha | Beta | Gamma | Delta | Epsilon | Lambda | Mu |
|---|---|---|---|---|---|---|
| Δ69/70 | L18F | L18F | T19R | S13I | G75V | T95I |
| Δ144 | D80A | T20N | G142D | W152C | T76I | Y144T |
| | D215G | P26S | E156G | | R246N | Y145SN |
| | R246I | D138Y | Δ157/158 | | Δ247-253 | |
| | | R190S | | | | |

**Figure S9 Sequence of the N-terminal domain in sarbecovirus and SARS-CoV-2 variants**

(A) Alignment of amino acid sequences of the NTD in sarvecoviruses analyzed in Figure 7E. Five loop regions are highlighted in red.

(B) List of NTD mutations in SARS-CoV-2 variants analyzed in Figure 7E.
